# N1-methylation of adenosine (m^1^A) in ND5 mRNA leads to complex I dysfunction in Alzheimer’s disease

**DOI:** 10.1101/2023.10.31.564907

**Authors:** Marko Jörg, Johanna E. Plehn, Marco Kristen, Marc Lander, Lukas Walz, Christine Lietz, Julie Wijns, Florian Pichot, Liliana Rojas-Charry, Katja M. Wirtz Martin, Nicolas Ruffini, Nastasja Kreim, Susanne Gerber, Yuri Motorin, Kristina Endres, Walter Rossmanith, Axel Methner, Mark Helm, Kristina Friedland

**Author notes:** To whom correspondence should be addressed., Correspondence may also be addressed to Mark Helm. The authors wish it to be known that, in their opinion, the first two authors should be regarded as joint first authors.

## Abstract

One mechanism of particular interest to regulate mRNA fate post-transcriptionally is mRNA modification. Especially the extent of m^1^A mRNA methylation is highly discussed due to methodological differences. However, one single m^1^A site in mitochondrial ND5 mRNA was unanimously reported by different groups. ND5 is a subunit of complex I of the respiratory chain. It is considered essential for the coupling of oxidation and proton transport. Here we demonstrate that this m^1^A site might be involved in the pathophysiology of Alzheimer’s disease (AD). One of the pathological hallmarks of this neurodegenerative disease is mitochondrial dysfunction, mainly induced by Amyloid β (Aβ). Aβ mainly disturbs functions of complex I and IV of the respiratory chain. However, the molecular mechanism of complex I dysfunction is still not fully understood. We found enhanced m^1^A methylation of ND5 mRNA in an AD cell model as well as in AD patients. Formation of this m^1^A methylation is catalyzed by increased TRMT10C protein levels, leading to translation repression of ND5. As a consequence, here demonstrated for the first time, TRMT10C induced m^1^A methylation of ND5 mRNA leads to mitochondrial dysfunction. Our findings suggest that this newly identified mechanism might be involved in Aβ-induced mitochondrial dysfunction.

## Introduction

RNA modifications are post-transcriptional changes to the chemical composition of ribonucleic acids. More than 170 different types of these alterations, which are thought to fine-tune RNA function (1–5) have been identified (6). They abundantly occur on transfer RNA (tRNA) and ribosomal RNA (rRNA). In contrast, occurrence, quantity and positioning of mRNA modification, especially methylation, are still highly discussed. One of these controversial mRNA methylations is m^1^A(7). Previous reports using commercially available antibodies mapped hundreds of m^1^A residues in eukaryotic RNA (8). These antibodies are directed against the modified nucleobase m^1^A. However, they were also found to bind cap structures (9, 10). A recent publication using a more stringent transcriptome-wide mapping approach for m^1^A RNA modification at single-base resolution only detected m^1^A in a low number of mRNAs (11). Irrespective of the approach used, one m^1^A modification coherently determined in all these different publications is the mitochondrial ND5 mRNA m^1^A at position 1374 (9, 11, 12). Biosynthesis of m^1^A is catalyzed by the writer methyltransferase TRMT10C (11). TRMT10C is the catalytic subunit of the mitochondrial methyltransferase that is responsible for introducing m^1^A and m^1^G at position 9 of human mitochondrial tRNAs ((mt)tRNAs) (13); it requires the (tetrameric) short-chain dehydrogenase/reductase SDR5C1 as a “structural” cofactor (13–15). In addition to acting as a methyltransferase, the TRMT10C-SDR5C1 complex together with the nuclease subunit PRORP constitutes the mitochondrial RNase P complex, which is responsible for the endonucleolytic release of tRNA-5’ ends from mitochondrial primary transcripts (14).

The protein encoded by the thus methylated mRNA is ND5, the fifth subunit of complex I of the respiratory chain. Complex I itself consist of 37 nuclear and 7 mitochondrially encoded subunits. It is essential for cellular energy production, providing 40% of the proton motive force required for ATP synthesis (13). Complex I is built up of a peripheral and a membrane-embedded arm (15). The mitochondrial-encoded subunit ND5 is located at the most distal position of the membrane arm. It provides a long transversal helix aligned parallel to the membrane arm. It was proposed that this helix might be used as a piston to transmit the energy released by the redox reaction in the peripheral arm to proton translocation in the membrane arm (16, 17). Therefore, ND5 is considered essential for the coupling of oxidation and proton transport (18).

Because of the location of the methylgroup in m^1^A on the Watson-Crick face, ND5 mRNA methylation on position 1374 within codon GCA is thought to disrupt base pairing during decoding on the mitoribosome, thereby impeding effective translation of ND5. This hypothesis was supported by polysome fractioning experiments suggesting repressed translation of m^1^A-consisting ND5 transcripts and m^1^A-induced ribosome stalling (11, 12). However, there is no direct experimental evidence that ND5 m^1^A mRNA methylation leads to reduced mitochondrial function or might even be involved in pathological alterations of mitochondrial function.

Mitochondrial dysfunction is one of the major pathologic hallmarks in sporadic Alzheimer’s disease (AD) (19–21), the most common form of AD (99 % of all AD cases). Age is the greatest known risk factor and mitochondria have long been implicated in the aging process and age-related diseases. Several alterations in mitochondrial function were associated with AD, such as reduced mitochondrial membrane potential, reduced ATP levels, enhanced reactive oxygen species (ROS) as well as impaired mitochondrial morphology, mitochondrial fusion and mitophagy (20, 22–26). These pathological changes are assumed to be strongly associated with reduced effectivity of the respiratory chain especially complex I (22, 27–30). Complex I is the major source of mitochondrial-derived ROS. Clinical investigations have reported impairment of complex I activity and reduced protein and mRNA levels of its nuclear-encoded subunits e.g. NDUFA2 or NDUFB3/7 in multiple zones of post-mortem AD brains as well as in blood cells such as lymphocytes or platelets (27). The findings are also recapitulated in cell and animal models which are characterized by increased levels of Amyloid β (Aβ) (31, 32). Data dealing with mitochondrial-encoded subunits of complex I such as ND5 are still missing. In recent AD genome-wide association studies, no single nucleotide polymorphisms (SNPs) in genes coding for complex I were identified, pointing to epigenetic or posttranscriptional mechanisms regulating complex I activity and expression (33, 34).

Therefore, we investigated the hypothesis that TRMT10C mediated ND5 mRNA methylation might contribute to complex I dysfunction in AD. Analyses were conducted on AD cell models as well as on post-mortem brain tissue, and supported by RNA-Seq data from AD patient studies. Investigations were based on a broad methodological spectrum ranging from western blot analysis, Illumina sequencing, RNA modification analysis and bioinformatics to mitochondrial respiration assays. We provide evidence that TRMT10C protein expression is enhanced in AD via a ROS-dependent measure leading to m^1^A ND5 mRNA methylation cumulating in reduced ND5 protein expression. Furthermore, we show that enhanced TRMT10C protein expression leads to impaired mitochondrial function, reflected in reduced mitochondrial respiration and membrane potential. With this study, we describe a completely new posttranscriptional mechanism how complex I protein expression and function is reduced in AD.

## Material and Methods

### Cell culture

The stably expressing APPwt HEK 293 cells and Control cells (Ctl) (untransfected HEK 293) were cultured as previously described (35, 36). The tetracycline-inducibaleTRMT10C (pTRMT10C) HEK cell line and control T-Rex-293 (Ctl) were cultured according to (37). For detailed information please see Supplementary Material.

### TRMT10C Knockdown

For siRNA-induced knockdown of TRMT10C, predesigned *Silencer*^TM^ select siRNA against TRMT10C (Thermo Fisher Scientific, 4392420, Assay ID: s29784) and the related *Silencer*^TM^ select siRNA negative control (Thermo Fisher Scientific, 4390843) were purchased. Transfection was conducted according to manufacturer’s protocol.

### Aβ oligomerization

Oligomerization of A**β**_1-42_ was performed according to manufacturer’s protocol. Aggregation was verified by Thioflavin T dye (AnaSpec, AS-88306) staining at 440Em/484Ex (38).

### Aβ_1-40_ Elisa

Measurement of A**β**_1-40_ levels was performed using an Amyloid beta 40 Human ELISA Kit (Invitrogen, KHB3482). For further information please see Supplementary Material.

### Western Blot

20-40 µg protein per lane was loaded on a 10% SDS polyacrylamide gel electrophoresis (PAGE). Western Blotting was conducted according to Heiser et al. (39). Information about antibodies is provided in Supplementary Materials.

### Real-Time quantitative PCR

qPCR was conducted according to manufacturer’s protocol. For detailed information please see Supplementary Materials.

### Generation of mitochondrial extracts

A mitochondria isolation was done to enrich the desired mitochondrial mRNA. Therefore HEK cells were washed with ice-cold PBS and seeded in large cell culture plates. After two days growing, medium was changed and on the third day the isolation of mitochondria was performed based on Sims *et al.* (40).

### Detecting misincorporation rate at position 1374 of ND5 mRNA

The method was adapted according to Safra et al. (11). Detailed information is provided in Supplementary Materials.

### Selective isolation and quantification of mitochondrial tRNAs from total HEK cell RNA

Detailed protocols how to selectively isolate and quantify mitochondrial tRNA are provided in Supplementary Material.

### Mitochondrial function assays

Mitochondrial membrane potential and mitochondrial respiration was measured according to (41). Details are described in Supplementary Materials.

### Human databases

Two different publicly available databases were used: RNA-Seq data from the Aging, Dementia and Traumatic Brain Injury (AgDemTBI) Study (primary publication: Miller J. A., *et al.* (42)) and the single cell (sc) RNAseq data from Mathys *et al.* (43). Details regarding the analysis of the data are provided in Supplementary Material.

### Human brain samples

Human brain material was provided via the rapid autopsy program of the Netherlands Brain Bank (NBB), which provides post-mortem specimens from clinically well documented and neuropathologically confirmed cases. Gyrus frontalis superior 3+4 was selected for controls as well as AD patients (more information in Table 1. and Supplementary Tables S4 and S5).

**Table 1:** Demographic data of AD subjects and healthy controls (more detailed overview in Supplementary Table S3.)

|  | Control | AD patients |
| --- | --- | --- |
| Number of samples | 12 | 13 |
| Male/Female | 6/6 | 5/8 |
| Age [years] (Mean ± SD) | 85.1 ± 9.0 | 83.9 ± 8.1 |
| Braak Stage (Mean ± SD) | 2.5 ± 1.0 | 4.7 ± 0.6 |

### Statistics

Data was analyzed with standard statistical methods performed in Graph Pad Prism (GraphPad Software, version 5.04, La Jolla, CA USA). Data are presented as the means ± SEM and are representatives of at least three independent experiments. Data points that were regarded as extreme statistical outliers were excluded using the ROUT and Grubb’s test.

### Figures

All figures were created with Biorender.com.

## Results

TRMT10C is posited installing m^1^A on mitochondrial ND5 mRNA. This mRNA methylation is liable to inhibit ND5 protein translation, a subunit of complex I of the respiratory chain (Figure 1A). Until now, it was unclear whether such a mechanism is impairing overall mitochondrial function, e.g. mitochondrial respiration and membrane potential, and if there is a connection to pathological alterations in neurodegenerative diseases such as AD. To investigate the role of TRMT10C in AD, we postulated the following experimentally verifiable hypotheses. These are: (i) TRMT10C protein levels are altered in AD cell model as well as in AD patients and (ii) as a consequence, m^1^A methylation of ND5 mRNA is increased, leading to (iii) reduced ND5 protein expression. Via this causal chain, enhanced m^1^A methylation of ND5 mRNA results in (iv) impaired mitochondrial function (Figure 1A).

**FIGURE 1.**
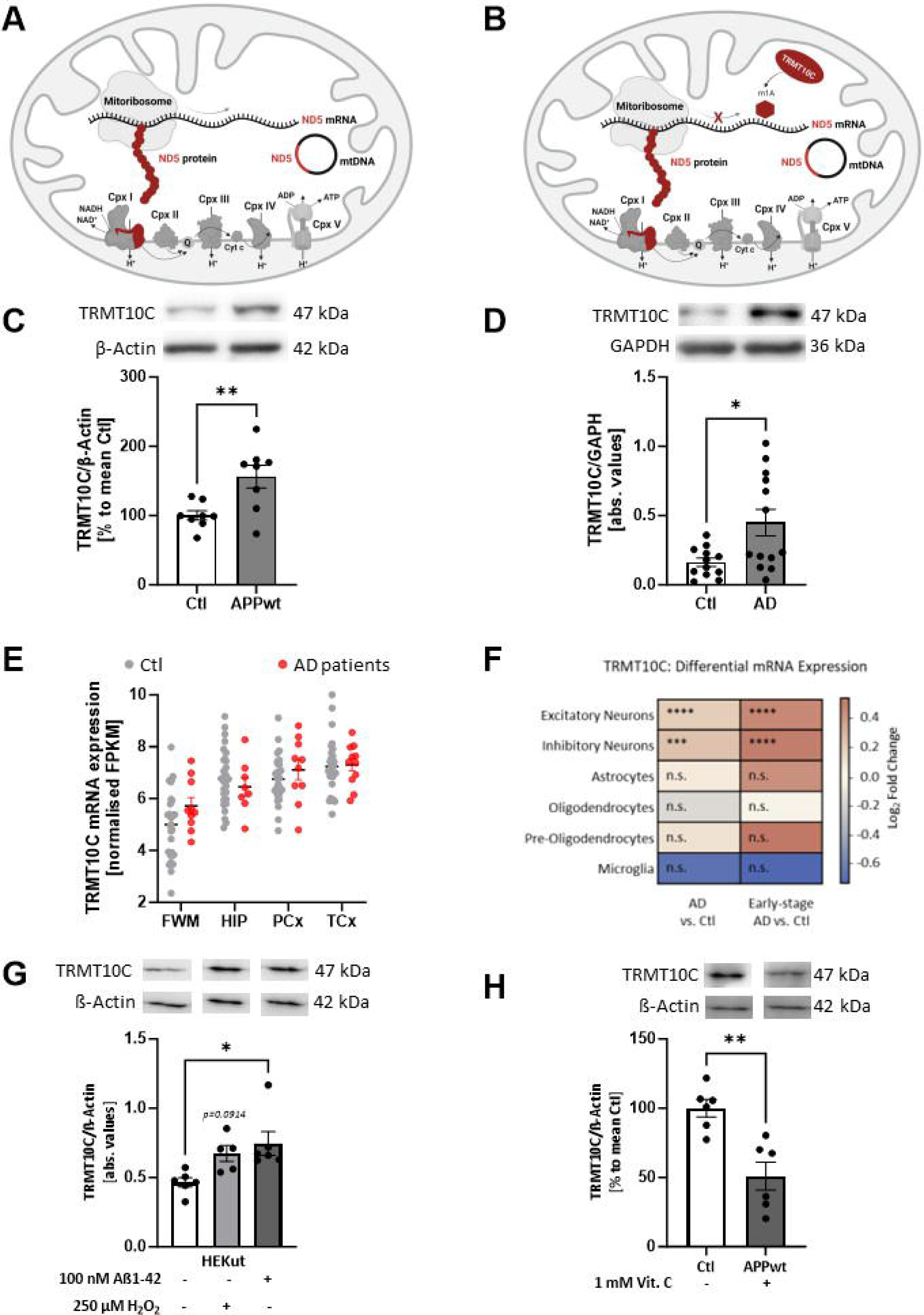
TRMT10C protein and mRNA levels are increased in AD cell and animal models and cortex samples of AD patients. A) Illustration of hypothesis and ND5 protein synthesis in mitochondria. m^1^A was found at position 1374 of the mitochondrial encoded ND5 mRNA transcript. m^1^A blocks Watson-Crick base pairing and leads to ribosome stalling, consequently the corresponding protein is translated inefficiently. It is assumed that decreased ND5 protein levels lead to complex I impairment. TRMT10C is the writer enzyme of this m^1^A site. B) Western Blot analysis shows a highly significant overexpression of TRMT10C in HEK APPwt cells compared to control cells. Mean ± SEM, n=8, unpaired t-test. C) Western Blot analysis of human frontal cortex samples (gyrus frontalis superior 3 + 4) from AD patients and aged-matched controls, Mean ± SEM, n=12 Ctl, n=13 AD, unpaired t-test. D) Comparison of TRMT10C mRNA expression in different brain regions of AD patients and healthy controls using the AgDemTBI database. TRMT10C mRNA is increased in all four investigated brain regions. WMPCx= White matter of Parietal Cortex, HIP= Hippocampus, PCx= Parietal Cortex, TCx= Temporal Cortex, FPKM= fragments per kilobase of exon model per million reads. Mean ± SEM, 29 controls (females 14, males 15) and 11 AD patients (females 2, male 8) were included into the analysis, unpaired t-tests (age: 77-100+). E) TRMT10C mRNA levels are significantly elevated in scRNA-Seq data from excitatory and inhibitory neurons, but unchanged in other brain cell types. This effect is particularly pronounced in Early-stage AD. 48 individuals (24 Ctl + 24 AD), 80 660 single-nucleus transcriptomes. F) Western blot of HEK-293 cells treated with 100 nM oligomeric Aβ_1-42_ for 24 hours showed significantly increased TRMT10C levels compared with corresponding control cells. H_2_O_2_ treatment (250 µM, 24 hours) tended to increase TRMT10C protein levels. Mean ± SEM, n=5-6, unpaired t-test. G) Western Blot analysis reveals significantly decreased TRMT10C protein levels in 1 mM Vitamin C (24 h) treated HEK APPwt cells in comparison to non-treated cells (HEK APPwt). Mean ± SEM, n=6, unpaired t-test. A)-G) *p<0.05, **p<0.01, ***p<0.001.

### Enhanced TRMT10C mRNA and protein expression in AD cell model and AD patients

To elucidate the first hypothesis, we used an AD cell model, AD patient post-mortem tissue and publicly available RNA-Seq databases. We investigated TRMT10C protein levels in HEK 293 cells stably overexpressing the wild-type human Amyloid precursor protein (APPwt), which displays a 10-fold increased Aβ_1-40_ level (22, 36). In this cell model TRMT10C protein levels were significantly increased compared to control cells (Figure 1B). In post-mortem frontal cortex samples from AD patients, received from The Netherlands Brain Bank (NBB), we detected significantly elevated TRMT10C protein levels compared to controls (Figure 1C, patient characteristics in Supplementary Table S4 and S5).

To further elaborate the role of TRMT10C in AD, we employed RNA-Seq data from two human databases (patient characteristics see Supplementary Table S4). One of these is the Aging, Dementia and Traumatic Brain Injury (AgDemTBI) Study (42, 44), in which the transcriptome of four different brain regions was analyzed. We compared FPKM values of all patients without TBI incident and confirmed AD diagnosis. TRMT10C mRNA levels were not altered in AD patients compared to controls (Figure 1D).

Using single-cell transcriptomic analysis (scRNA-Seq) examining 6 major brain cell types from the frontal cortex of 24 AD patients and 24 age-matched controls, we observed increased TRMT10C mRNA levels in excitatory and inhibitory neurons. In astrocytes, oligodendrocytes, precursor-oligodendrocytes and microglia no significant changes were found (Figure 1E) (43). This neuron-specific increase was already significant in early stages of AD and still present in later stages.

### Aβ mediated oxidative stress enhances TRMT10C protein expression

Our data so far using an Aβ overexpressing HEK APPwt cell model, NBB AD postmortem tissue and databases, suggested that Aβ might play a role in TRMT10C overexpression. However, other pathological changes e.g. mitochondrial dysfunction associated with elevated oxidative stress might also contribute to these alterations. To follow this idea, HEK Ctl cells were treated with oligomeric Aβ_1-42_ (100 nM) or H_2_O_2_ (250 µM) for 24 h and TRMT10C protein expression was analyzed. Under both stressors, we observed elevated TRMT10C protein expression (Figure 1F). A well described mechanism how elevated Aß levels impair mitochondrial function, is the increase of oxidative stress generated via complex I dysfunction (22). Therefore, we scavenged reactive oxidant species using Vitamin C (1000 µM). This treatment results in the rescue of TRMT10C protein expression in HEK APPwt cells (Figure 1G) suggesting that oxidative stress plays a vital role in the newly described alterations in our AD cell model.

### Enzymatic partner subunits of TRMT10C are differently affected in AD

TRMT10C requires the cofactor SDR5C1 to exhibit methyltransferase activity on position 9 of (mt)tRNAs (13); whether this is also the case for the methylation of ND5 A1374 has not been investigated yet (Figure 2A). The TRMT10C-SDR5C1 complex is also an essential part of mtRNase P, required for cleavage by the actual nuclease subunit PRORP (14). In the light of our findings on TRMT10C, this raised the question, whether the expression of SDR5C1 and PRORP is also altered in AD. Surprisingly, PRORP protein levels were significantly decreased in HEK APPwt cells (Figure 2B). In AD frontal cortical post-mortem samples, no significant changes in PRORP protein expression were detected (Figure 2C). Likewise, no differences were found on mRNA levels in both RNA-Seq data sets (Figure 2D for PRORP data are only available in the AgDemTBI Study but not in Mathys *et al.* study.) .

**FIGURE 2.**
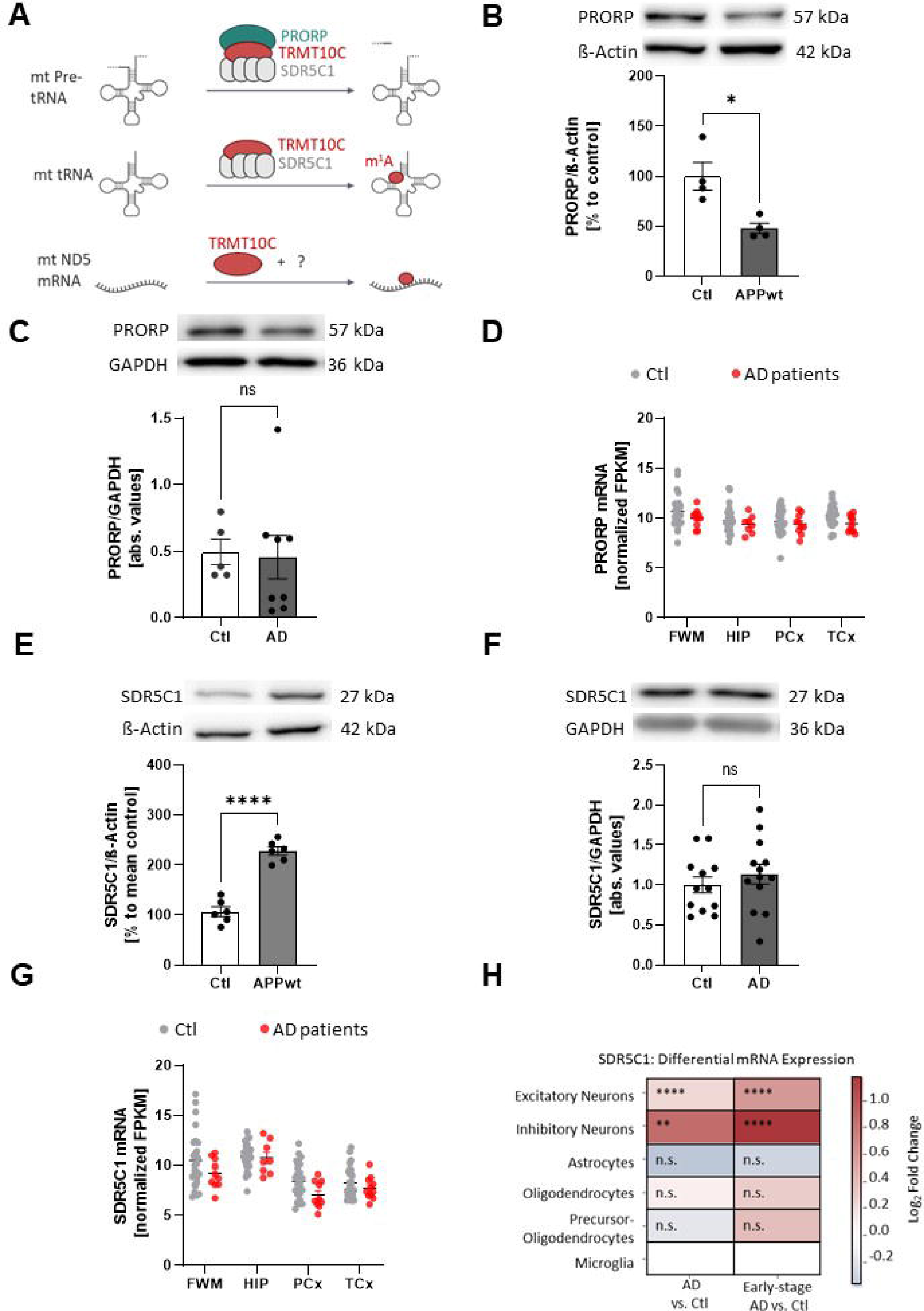
Western Blot analysis and evaluation of human databases reveal different behavior of mitochondrial RNase P subunits SDR5C1 and PRORP. A) Schematic overview about the three different functions of TRMT10C. First, a tripartite complex, constituted of PRORP, TRMT10C and SDR5C1 cleaves mitochondrial tRNAs out of the polycistronic transcript. Second, TRMT10C methylates mitochondrial tRNAs in combination with an SDR5C1 tetramer. Third, TRMT10C methylates ND5 mRNA at position 1374. B) Western Blot analysis indicates significantly reduced PRORP levels in HEK APPwt cells, n=4, Mean ± SEM, unpaired t-test. C) Western Blot analysis of human frontal cortex samples from AD patients and aged-matched controls shows no significant changes in PRORP protein levels, Mean ± SEM, n=12 Ctl, n=13 AD, unpaired t-test. D) Comparison of PRORP mRNA expression in different brain regions of AD patients using the “Aging, Dementia & TBI Study” database. No significant changes were detected in all four brain regions. Mean ± SEM, n=13-15 Ctl n=5-8 AD, unpaired t-tests (age: 77-100+, sex: male), WMPCx= White matter of Parietal Cortex, HIP= Hippocampus, PCx= Parietal Cortex, TCx= Temporal Cortex, FPKM= fragments per kilobase of exon model per million reads. E) Western Blot of HEK APPwt cells shows significantly elevated SDR5C1 levels in comparison to Ctl. Mean ± SEM, n=6, unpaired t-test. F) In Western Blot analysis of human frontal cortex samples from AD patients and aged-matched controls no alterations in SDR5C1 protein levels can be found, Mean ± SEM, n=12 Ctl, n=13 AD, unpaired t-test. G) Comparison of SDR5C1 mRNA expression in four different brain regions of AD patients using the AgDemTBI database. SDR5C1 mRNA is no significantly altered in AD patients. Mean ± SEM, 29 controls (females 14, males 15) and 11 AD patients (females 2, male 8) were included into the analysis, unpaired t-tests (age: 77-100+), WMPCx= White matter of Parietal Cortex, HIP= Hippocampus, PCx= Parietal Cortex, TCx= Temporal Cortex, FPKM= fragments per kilobase of exon model per million reads. H) SDR5C1 mRNA levels are significantly elevated in scRNA-Seq data from excitatory and inhibitory neurons, but unchanged in other brain cell types. This effect is particularly pronounced in Early-stage AD. 48 individuals (24 Ctl + 24 AD), 80 660 single-nucleus transcriptomes. A)-H) *p<0.05, **p<0.01, ***p<0.001.

Next, we focused on SDR5C1 and found significantly enhanced protein levels in HEK APPwt cells but not in human post-mortem tissue (Figure 2E, F). The AgDemTBI Study database showed no significant changes in SDR5C1 mRNA expression (Figure 2G). The scRNA-Seq data revealed a strong upregulation of SDR5C1 mRNA levels only in excitatory and inhibitory neurons (Figure 2H).

### m^1^A levels are increased in AD cell model and patients

After showing that the m^1^A writer enzyme TRMT10C was consistently upregulated in AD models and human AD cortex samples, we scrutinized a plausible implication, namely if m^1^A levels in ND5 mRNA would be elevated in AD pathology as pointed out in our hypothesis (ii). We first analyzed our AD cell model using a site-specific analysis method based on m^1^A-induced misincorporation during reverse transcription and Illumina sequencing (workflow in Supplemental Figure S1). Knowing that yields of misincorporation and reverse transcription arrest both depend heavily on reaction conditions and the reverse transcriptase, we used SuperScript IV (SS-IV) as it provides high read-through capability on modification sites, thus yielding more full-length product suitable for RT-PCR and mismatch rate analysis (45).

We found elevated misincorporation levels at position 1374 of ND5 mRNA in total RNA samples from HEK APPwt cells in comparison to Ctl (Figure 3A), as well as in RNA isolated from mitochondrial extracts (Figure 3B). In both sets mismatch was mainly composed of T and G, suggesting that m^1^A mainly evokes incorporation of A and C during RT with SS-IV. In addition, increased ND5 mRNA methylation was reflected in elevated jump rates of HEK APPwt in both, total RNA and mitochondrial RNA preparations (Figure 3A). Jumps are nucleotide skipping events that occur due to the disruption of the reverse transcriptase by m^1^A and represent a second, independent parameter to assess m^1^A methylation (46). In our NBB samples, after a first analysis m^1^A mismatch rate was not altered. This picture changed after we separated the data according to the Braak stages, a measure for the neuropathologic stadium of all patients (47). A significant increase in mismatch and jump rates was revealed when comparing AD with Braak stage 5+6 versus controls with Braak stage 0-3 (Figure 3C). 2 AD patients carried a G13708A SNP, which is prevalent in the Eurasian J haplogroup. This SNP was shown to prevent methylation of m^1^A two bases downstream (position 1374 in ND5 mRNA equates position 13710 of mtDNA) (11). In these patients, we obtained no mismatch at all. Therefore, these individuals were omitted from the analysis.

**FIGURE 3.**
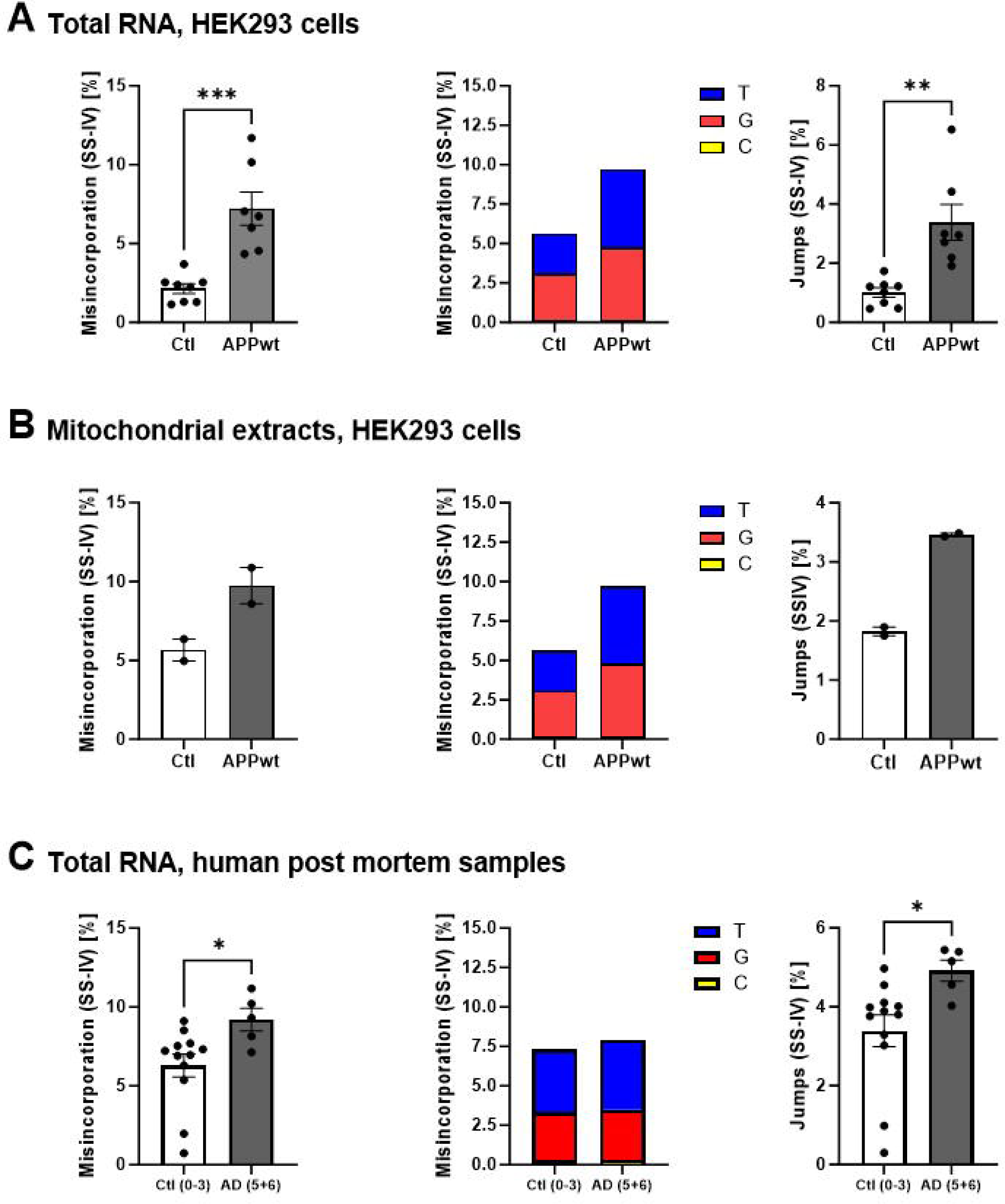
m^1^A levels at position 1374 in ND5 mRNA are increased in AD model cells and in primary visual cortex samples from human patients. A) m^1^A-induced misincorporation and jumps are significantly elevated in APPwt cells in comparison to Ctl. Shown are single replicate values, misincorporation split into individual bases and jump rate. Mean ± SEM, n=7-8, unpaired t-test. B) RNA isolated from mitochondrial extracts confirms enhanced m^1^A methylation in HEK APPwt cells in comparison to Ctl. Mean ± SEM, n=2. C) m^1^A-induced misincorporation and jumps are significantly increased in human AD postmortem brain samples compared to non-demented controls. Mean ± SEM, n=5-12, unpaired t-test. A)-C) For every data set sequencing reads at position 1374 of mt ND5 mRNA are shown as misincorporation including different biological replicates, misincorporation split into individual bases and jumprate. *p<0.05, **p<0.01

Biochemical evidence shows that TRMT10C and SDR5C1 form a stable sub-complex that is active as a tRNA-methyltransferase (14) and is uniquely able to methylate both adenosine and guanine nucleotides at position 9 of (mt)tRNA. 19 of the 22 mt-tRNAs contain either A or G at position. Hence, an overexpression of TRM10C might be expected to increase overall methylation levels in (mt)tRNA. Therefore, we isolated mitochondrial tRNA from HEK Ctl and HEK APPwt cells and quantified m^1^A and m^1^G levels. We observed no significant alterations (Supplemental Figure S2A and B), indicating either, that mt tRNAs were already fully modified, or that excess TRM10C, outside a complex with SDR5C1, does not target (mt)tRNAs for methylation.

### ND5 protein levels are decreased in AD cell model and brains of human patients

Since m^1^A ND5 mRNA methylation is hypothesized to impede Watson-Crick base pairing, (mt)tRNA would fail to bind to the ND5 mRNA codon during the translation process. Defective or absent binding would result in abortive or at least in a disruption of ND5 protein synthesis. However, the direct correlation between m^1^A methylation and ND5 protein levels has not been studied so far. For this reason, we performed Western Blots of ND5 in our AD cell model and AD patients’ samples examined to verify our hypothesis (iii).

In HEK APPwt cells, ND5 protein levels were significantly decreased in comparison to control cells (Figure 4A), suggesting a disruptive nature of m^1^A on ND5 protein biosynthesis. Again, in the NBB samples, no significant changes in ND5 protein levels were detected (Figure 4B). However, similar to our mismatch results, we observed significantly reduced protein expression of ND5 when comparing AD patients with Braak stage 5+6 versus controls with Braak stage 0-3 (Figure 4B and C). To investigate if the altered protein levels of ND5 might be due to reduced ND5 mRNA levels, we conducted RT-qPCR in our AD cell model and postmortem tissue. No difference in the ND5 mRNA level of HEK APPwt cells compared to the controls were observed (Fig.4D). In the post-mortem tissue, we even found increased ND5 mRNA levels in all AD patients. Similar effects were detected in AD patients with Braak stage 5+6 versus controls with Braak stage 0-3 (Fig. 4E and F). The quantification was performed using a custom ND5 primer which omits the m^1^A site during hybridization.

**FIGURE 4.**
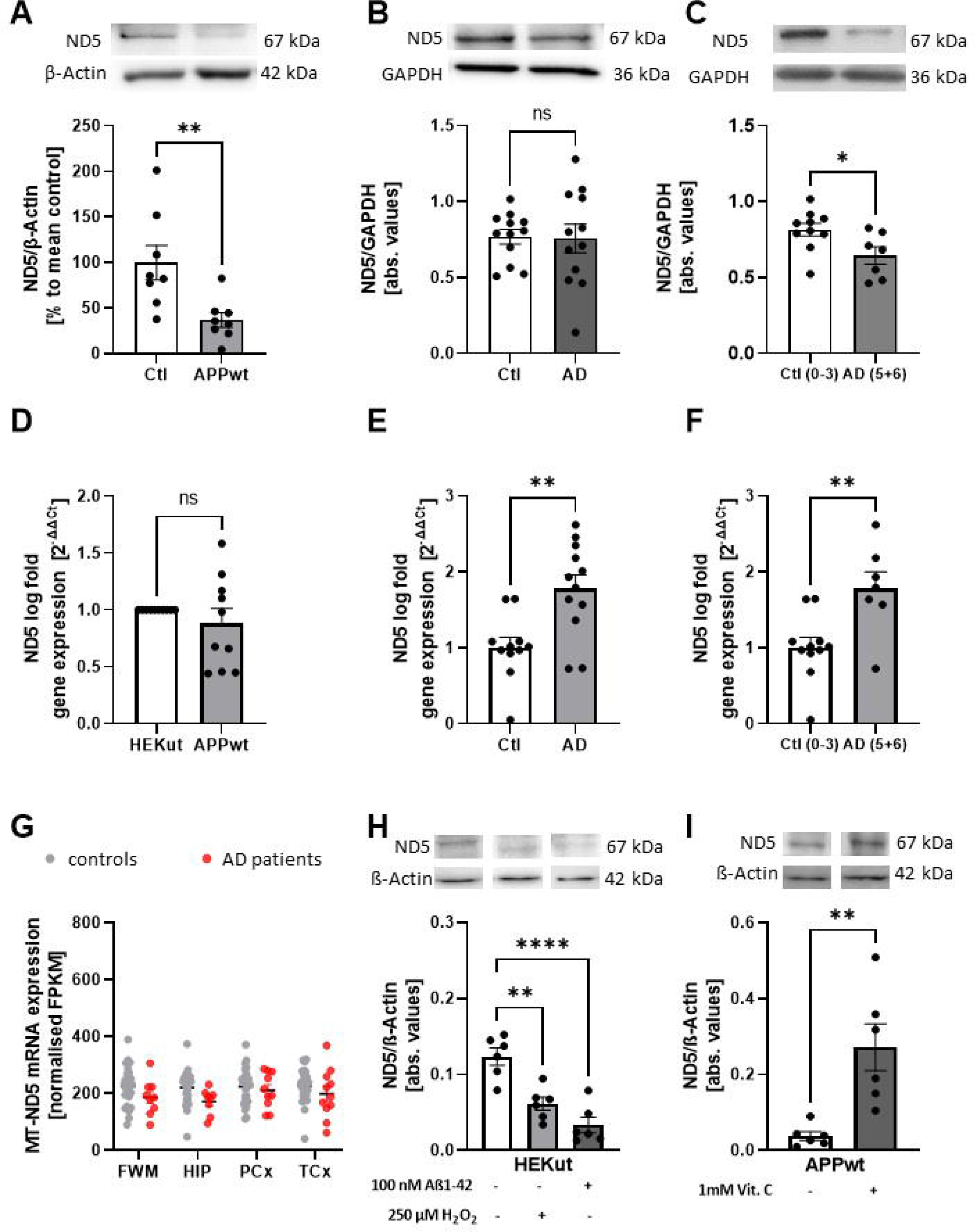
ND5 protein levels are decreased in AD cell and animal models and in AD patients brains. A) Western Blot analysis reveals reduced ND5 protein levels in HEK APPwt cells. Mean ± SEM, n=8, unpaired t-test. B) ND5 protein levels are not altered between frontal cortex samples from AD patients and aged-matched controls, Mean ± SEM, n= 12 Ctl, n=13 AD, unpaired t-test. C) Comparison between AD patients at Braak stage 5+6 and Ctl at Braak stage 0-3 reveals significantly reduced ND5 levels Mean ± SEM. D) ND5 mRNA expression in HEK APPwt cells not different to HEK Ctl cells. Determined using qPCR, Mean ± SEM, n=10, unpaired t-test. E) ND5 mRNA levels increased in frontal cortex samples from AD patients and aged-matched controls using RT-qPCR, Mean ± SEM, n= 12 Ctl, n=13 AD, unpaired t-test. F) Comparison between AD patients at Braak stage 5+6 and Ctl at Braak stage 0-3 still shows increased ND5 mRNA levels Mean ± SEM. G) No changes in ND5 mRNA expression in parietal cortex samples using AgDemTBI RNAseq data from human AD patients in comparison to healthy controls. Mean ± SEM, 29 controls (females 14, males 15) and 11 AD patients (females 2, male 8) were included into the analysis, unpaired t-test (age: 77-100+, sex: male), WMPCx= White matter of Parietal Cortex, HIP= Hippocampus, PCx= Parietal Cortex, TCx= Temporal Cortex, FPKM= fragments per kilobase of exon model per million reads. I) Protein levels of mt-ND5 in HEK-293 cells treated with 100 nM oligomeric Aβ_1-42_ and 250 µM H_2_O_2_ for 24 hours were significantly decreased compared to the untreated control. Mean ± SEM, n=6, unpaired t-test. I) Protein levels of mt-ND5 in Vitamin C treated HEK APPwt cells were significantly increased compared to non-treated HEK APPwt cells. Mean ± SEM, n=6. A)-F) *p<0.05, **p<0.01, ***p<0.001.

To further assess the status of ND5 in human patients, we used data from the AgDemTBI study and observed no significant differences between AD patients and healthy controls (Figure 4G). Unfortunately, no information about ND5 mRNA was available in the scRNA-Seq study from Mathys *et al.* (43).

To check whether changes in ND5 protein expression are also mediated by elevated Aβ_1-42_ levels and enhanced oxidative stress, we treated HEK Ctl cells with Aβ_1-42_ (100 nM) and H_2_O_2_ (250 µM) for 24 h. ND5 protein levels were significantly reduced under both stressors (Figure 4H). To evaluate if Aβ induced oxidative stress is involved in reduced ND5 protein expression in HEK APPwt cells, Vitamin C (1000 µM) was used again as an antioxidant. ND5 protein expression was rescued under this condition (Figure 4I).

To exclude that Aß might impair the translation of all mitochondrial encoded subunits of complex I by nonspecific effects, ND1 protein expression was examined in HEK APPwt cells. We found no significant differences in ND1 protein expression between HEK Ctl cells and HEK APPwt cells (Supplemental Figure S3).

### TRMT10C mediated m^1^A mRNA methylation correlates with repressed ND5 protein translation

Although TRMT10C was upregulated, m^1^A methylation of ND5 mRNA was increased and ND5 protein levels downregulated in several AD model systems and human patients, a postulated causality between these findings needed to be further substantiated. For this reason, we used the tetracycline-inducible HEK293 cell system overexpressing TRMT10C (pTRMT10C) (37) to scrutinize the hypothesis (iv). We first correlated the effects of 0.1 µg/mL and 1 µg/mL tetracycline after 24 h on TRMT10C protein expression and on m^1^A ND5 mRNA methylation (Figure 5A). To exclude effects of tetracycline itself (48, 49), we also investigated the effects of 0.1 µg/mL and 1 µg/mL tetracycline in the corresponding control cell line (Ctl). A concentration dependent effect on TRMT10C protein expression was observed in pTRMT10C cells, which, importantly, also correlated with m^1^A mRNA methylation (Figure 5B).

**FIGURE 5.**
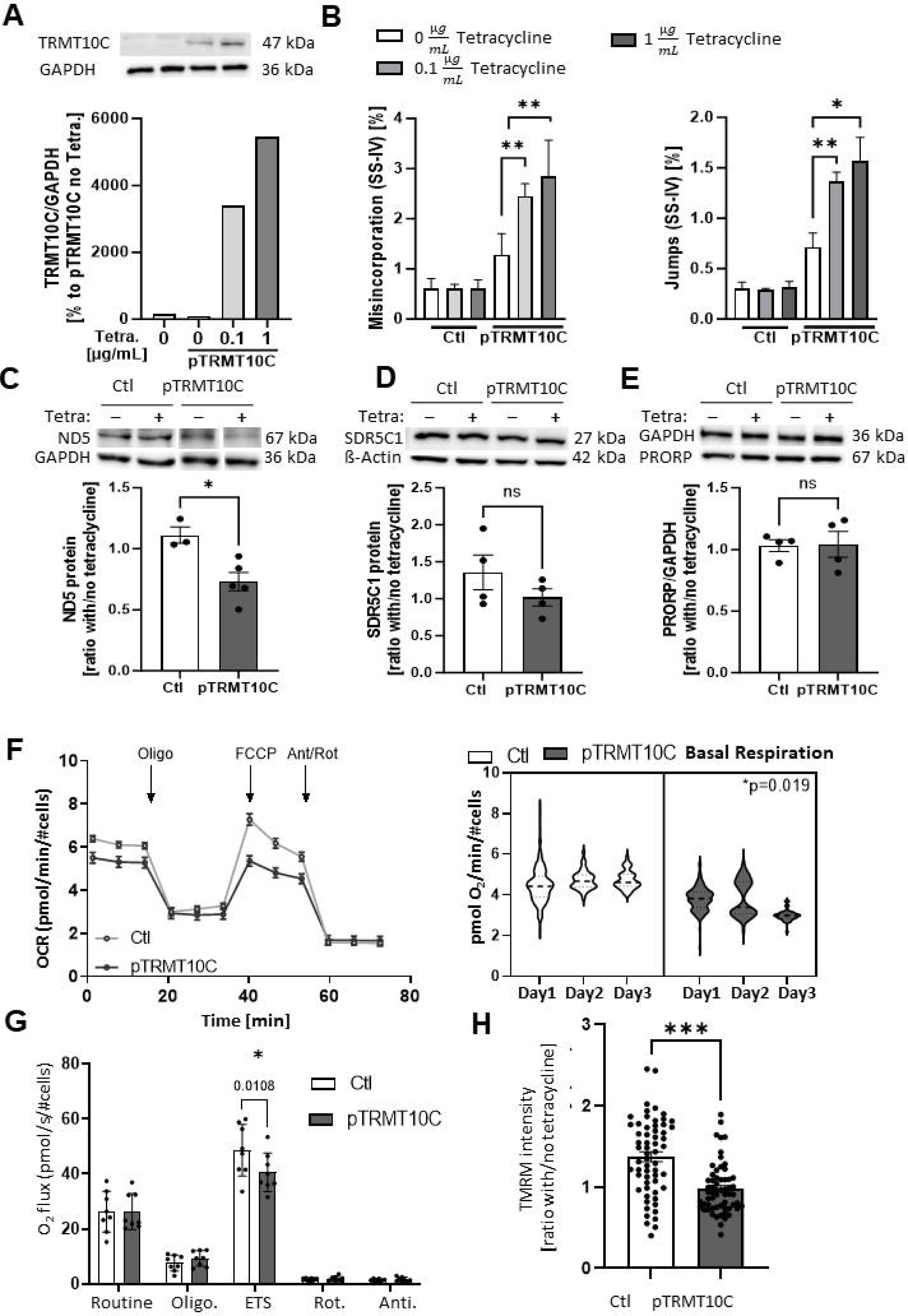
Tetracycline-induced overexpression of TRMT10C and its effects on m^1^A, ND5, SDR5C1, PRORP protein levels and mitochondrial function. A) Western Blot of pTRMT10C cells and control cell line after incubation with different tetracycline concentrations. n=1. B) Misincorporation and jump rate at position 1374 of mitochondrial ND5 mRNA after RT-PCR with SSIV, Mean + SEM, n=4, unpaired t-test. C) Western Blot analysis reveals significantly reduced ND5 protein levels in pTRMT10C cells after addition of tetracycline in comparison to Ctl. Mean ± SEM, n=3 Ctl, n=5 pTRMT10C, unpaired t-test. D) SDR5C1 expression is unchanged in Western Blots of pTRMT10C cells and Ctl after addition of tetracycline. Mean + SEM, n=4. E) Western Blot analysis shows no changes of PRORP expression in pTRMT10c cells and Ctl after addition of tetracycline. Mean + SEM, n=4. A)-E) unpaired t-test *p<0.05, **p<0.01. F) Representative oxygen consumption curve of Ctl and pTRMT10C cells, measured in the XF96 Extracellular Flux analyzer. Violine plots for Basal Respiration are shown from three individual experiments. n=3, nested analysis. G) Oxygen consumption after addition of different chemical compounds in the roboros oxygen electrode. Oligo. = Oligomycin, ETS= Electron Transport System, Rot.= Rotenone, Anti.=Antimycin, Mean ± SD, n=8. H) TMRM staining reveals that mitochondrial membrane potential is significantly decreased in pTRMT10C cells after addition of 1 µg/mL tetracycline in comparison to Ctl, Mean ± SEM, data points represent values of every individual well from n=4 different days. Unpaired t-test.

To determine whether increased mRNA methylation results in reduced ND5 protein synthesis, we used 1 µg/mL tetracycline for 24 h to induce a robust TRMT10C protein overexpression and corresponding m^1^A methylation of ND5. ND5 protein levels were significantly decreased in pTRMT10C cells in comparison to control cells incubated with tetracycline (Figure 5C). Consistent with the analysis described above, SDR5C1 and PRORP protein expression was not affected by TRMT10C overexpression (Figure 5D and E) neither was ND5 mRNA expression (data not shown). Additionally, we found that increased TRMT10C alone does not lead to any change in (mt)tRNA levels, given that after incubation of pTRMT10C cells with 1 µg/mL tetracycline, the ratio between mitochondrial and cytosolic tRNAs remained unchanged (see Supplementary Figure S4A). In addition, no alteration in m^1^A and m^1^G levels in (mt)tRNA were observed (see Supplementary Figure S4B and C).

### TRMT10C overexpression induces severe mitochondrial deficits

Because TRMT10C-induced m^1^A methylation in ND5 mRNA lowers ND5 protein levels, we next assessed whether this reduced ND5 protein expression would be sufficient to induce mitochondrial dysfunction. Therefore, we examined mitochondrial function after tetracycline-induced induction of TRMT10C (37).

We first used a Seahorse XFe 96 Extracellular Flux Analyzer and compared the oxygen consumption rates (OCR) to control cells (50). Oligomycin was first injected to inhibit the ATP synthase, impacting ATP production. The following injection was FCCP to evaluate the maximal respiratory capacity through uncoupling of oxygen uptake from ATP production. Finally, rotenone and antimycin were injected to inhibit complex I and III to estimate non-mitochondrial respiration. Representative graphs are shown in Figure 5F. Remarkably, basal respiration was significantly reduced in pTRMT10C cells as compared to control cells also treated with tetracycline (Figure 5F, Supplementary Figure S5). To further verify our findings, we performed high-resolution respirometry using an OROBOROS Oxygraph-2k, which confirmed a decreased electron transport system capacity in pTRMT10C cells incubated with tetracycline in comparison to Ctl (Figure 5G). In line with this, the mitochondrial membrane potential quantified by TMRM staining was reduced in TRMT10C-overexpressing cells (Figure 5H).

### Loss-of-function experiments validate the causal chain from TRMT10C to ND5 protein expression

Our previous experiments established the effects of increased TRMT10C expression on mRNA methylation, ND5 levels, and respiration. To support the hypothesis of a molecular mechanism causally connecting these observations, we lowered TRMT10C levels using RNAi.

TRMT10C protein expression was strongly reduced by siRNA to 30 % compared to the scrambled negative control and untreated HEK APPwt cells (Figure 6A). This knockdown leads to a reduction of m^1^A induced mismatch and jump rates (Figure 6B and C). ND5 protein expression was significantly restored (Figure 6D). We observed no changes in Aβ_1-40_ levels (Figure 6E and F).

**FIGURE 6.**
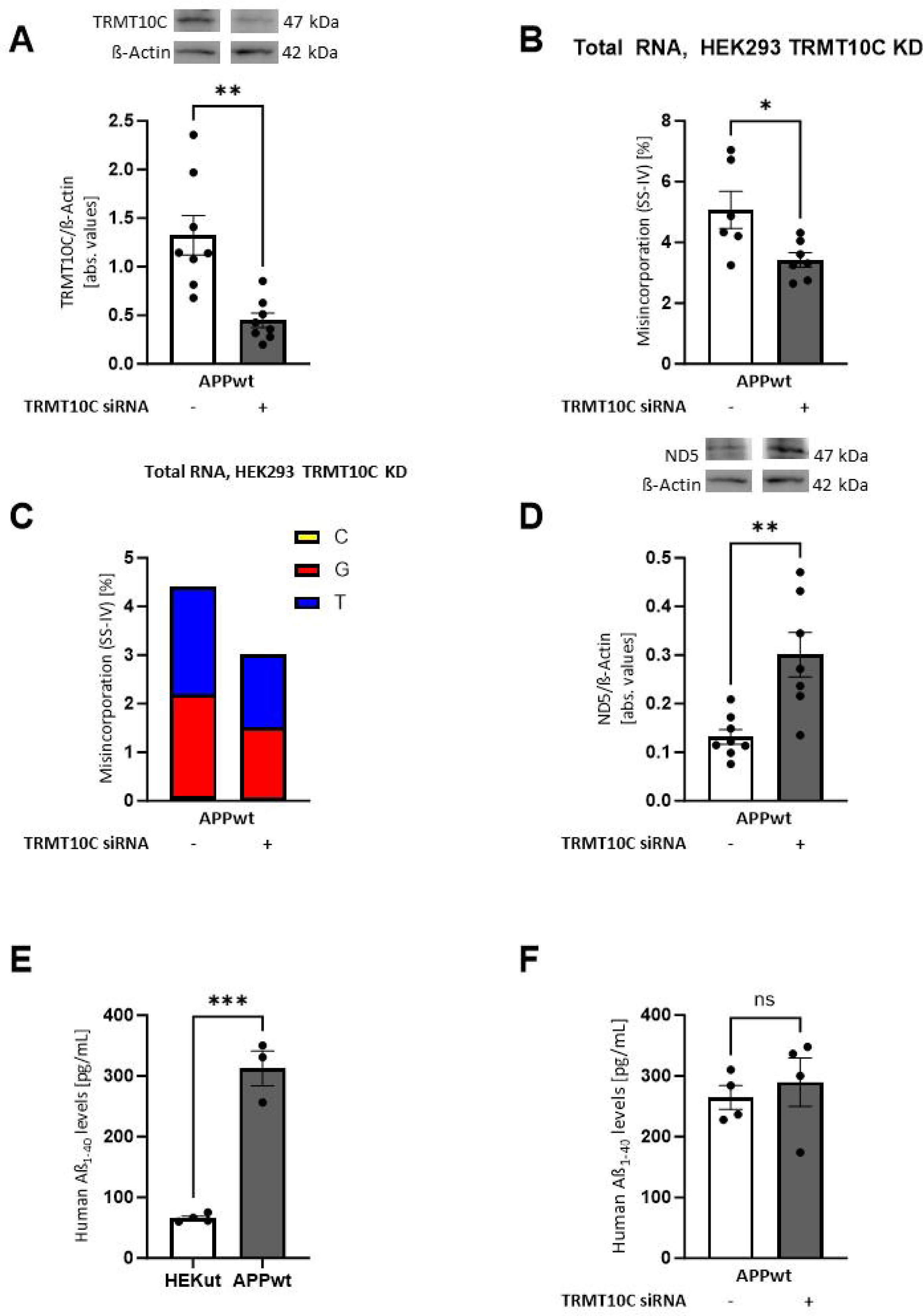
siRNA induced knockdown of TRMT10C and its effects on m^1^A mismatch and jump rate and mt-ND5 protein levels. A) Western Blot of siRNA induced TRMT10C KD cells. Mean ± SEM, n=8, unpaired t-test. B), C) Misincorporation rate at position 1374 of mitochondrial ND5 mRNA after RT-PCR with SS-IV after KD siRNA induced KD of TRMT10C. For every data set sequencing reads at position 1374 of mt ND5 mRNA are shown as misincorporation including different biological replicates. Mean ± SEM, n=6-7, unpaired t-test. D) Western Blot analysis reveals significantly increased ND5 protein levels in siRNA induced TRMT10C KD cells in comparison to non-KD cells. Mean ± SEM, n=3, unpaired t-test. A)-D) *p<0.05, **p<0.01, ***p<0.001.

## Discussion

The role of TRMT10C-induced m^1^A methylation of ND5 mRNA for mitochondrial function under physiological conditions and in the context of neurodegenerative diseases such as AD has not been investigated yet. Here, we provide first evidence that TRMT10C protein levels are increased in an AD cell model and in post-mortem tissue and RNA-Seq data from AD patients, plausibly leading to increased m^1^A methylation of ND5 mRNA. Increased levels of methylated ND5 mRNA correlate with reduced ND5 protein levels in AD models and patients. Similarly, we demonstrate that TRMT10C overexpression in HEK293 cells results in increased m^1^A methylation of ND5 and reduced ND5 protein levels. These results suggest that TRMT10C mediated m^1^A methylation of ND5 mRNA might be an important factor to control mitochondrial function and to trigger complex I dysfunction in AD.

TRMT10C is part of a multifunctional protein complex, playing a role in RNA processing, tRNA and mRNA methylation. TRMT10C and SDR5C1 form a stable complex with methyltransferase activity. Together with an additional subunit, PRORP, they form the mtRNase P with pre-tRNA processing activity. For ND5 m^1^A mRNA methylation, Safra *et al.* showed that TRMT10C is required, yet neither the involvement of SDR5C1 nor PRORP in this process was investigated. Here, we show that the protein and mRNA levels of TRMT10C and the other subunits of mtRNase P mainly PRORP are not affected similarly in AD. With regard to our data from pTRMT10C cells, we assume that overexpression of TRMT10C alone does not alter mtRNase P activity in AD. These results are supported by findings that the deleterious effect of Aβ on mitochondrial function are not explained by a specific inhibition of mtRNase P or its tRNA methyltransferase subcomplex (51).

Since the TRMT10C and SDR5C1 sub-complex also methylate (mt)tRNA, we asked whether the increased protein expression of both subunits might have an impact on (mt)tRNA methylation thereby regulating mitochondrial translation. We found no significant differences in m^1^A and m^1^G (mt)tRNA levels. There are two possible explanations for these findings. First, (mt)tRNA might already be fully methylated and an increase in TRMT10C and SDR5C1 expression cannot further increase methylation levels. Second, for tRNA modification TRMT10C and SDR5C1 need to form a stable subcomplex active as a methyltransferase (52). In AD however, Aß binds to SDR5C1 (53) and might thereby impair the formation of the stable subcomplex needed for (mt)tRNA interaction. Elevated TRMT10C levels might then be available to methylate ND5 mRNA. However, our results suggest that the observed changes in ND5 protein levels are not induced by altered mt(tRNA) mediated changes in protein synthesis.

The m^1^A methylation level in ND5 mRNA at position 1374 is increased in our AD cell model as well as in human AD patients. The Watson-Crick disruptive nature of m^1^A is hypothesized to prevent effective translation of the modified codons. This assumption can be applied with confidence for m^1^A sites residing in the first two nucleotides of the codon triplet, since Watson-Crick paring at these sites are known to be rigidly monitored on translating ribosomes. However, the case at hand features m^1^A1374 at the so-called “wobble” position of the codon, where non-standard base pairings are very frequent. As the mitochondrial translation apparatus significantly deviates from canonical ones, a productive non-standard base pairing, e.g. involving the m^1^A-Hoogsteen edge, remains possible. However, in HEK cells it was demonstrated that m^1^A within in a lysine codon (AAA), did not allow protein synthesis at all (54). This distinct inhibition was observed independent of the location of m^1^A within the codon. Even at the wobble position, which is usually less restrictive in terms of base pairing geometry, the methylation hindered protein synthesis (54).

Beside Aβ, Li *et al.* demonstrated that other mitochondrial stressors such as hypoxia or MitoBloCK-6 inhibiting protein import into mitochondria lead to increased m^1^A levels in ND5 mRNA. Furthermore, m^1^A methylation of ND5 mRNA seems to be tissue-specific with high levels in brain and blood and low levels in heart and muscle (11). This tissue-specific misincorporation profile of ND5 mRNA might be one piece in the puzzle why mitochondrial function is mainly impaired in brain and blood cells of AD patients and not in muscle or heart. Another study demonstrated that m^1^A mRNA levels are upregulated in AD patients and that oxygen-glucose deprivation/reoxygenation in primary cortical mouse neurons leads to m^1^A peaks without deciphering the exact position of the methylation.

ND5 protein levels are downregulated in our AD cell model and post mortem brain samples. Several lines of evidence demonstrate that complex I dysfunction is one central malfunction in AD (27). Holper *et al.* (27) observed downregulated complex I in different regions of the cortex, as well as in the hippocampus of AD patients. The protein expression of the mitochondrial encoded subunit ND5 in AD was never investigated until now. However, protein levels of several nuclear encoded complex I subunits were found to be decreased in AD patient post-mortem tissue (25). We established a first connection between TRMT10C induced m^1^A ND5 mRNA methylation resulting in reduced ND5 protein expression and mitochondrial dysfunction. TRMT10C overexpression resulted in elevated m^1^A ND5 mRNA methylation, reduced basal mitochondrial respiration and mitochondrial membrane potential. As mentioned above SDR5C1 and TRMT10C work in concert as a complex to install tRNA methylation. However, we observed no altered SDR5C1 or PRORP protein levels, the third subunit of mtRNAase P, in this cell model. These results can be explained by three hypothetical models: First, TRMT10C acts as a monomer to methylate ND5 mRNA. Second, SDR5C1 protein is abundantly expressed and forms a subcomplex to methylate mRNA. Third, another, yet unknown partner, might interact with TRMT10C. Keeping in mind that a complex of TRMT10C and SDR5C1 is required to methylate tRNA, the second hypothesis might be favorable.

In contrast, when we knock down TRMT10C protein expression in our HEK APPwt cell model, m^1^A mismatch rate is significantly reduced and ND5 protein levels are rescued. We also investigated Aβ_1-40_ levels and found no significant changes.

Finally, we provide mechanistic data showing that both Aβ_1-42_ and oxidative stress lead to enhanced TRMT10C and reduced ND5 protein expression. The pathophysiology of AD is driven by its hallmark oligomeric Aβ_1-42_ but also by other major dysfunction such as mitochondrial energy failure accompanied by elevated oxidative stress. Both, Aβ_1-42_ and oxidative stress mediated changes in TRMT10C and ND5 protein expression are reverted scavenging reactive oxidant species using Vitamin C (1000 µM). We previously demonstrated that this treatment results in reduced oxidative stress and lowered Aβ_1-40_ burden in HEK APPwt cells (22). These changes were accompanied by a reduction in BACE1 activity (22). BACE1 regulates, together with ADAM and γ-secretase complex, APP processing (55, 56).

Together, these results prompt us to conclude that m^1^A methylation of ND5 mRNA might fine-tune mitochondrial function under physiological conditions to the energetic needs of the cells (11). Under pathophysiological conjectures e.g. of enhanced mitochondrially generated ROS or elevated Aβ levels crossing the threshold of healthy aging toward pathological aging and AD, TRMT10C protein expression and m^1^A methylation levels of ND5 mRNA are strongly elevated resulting in repression of ND5 protein translation. Together with the Aβ induced inhibition of protein transport into mitochondria, reduced mRNA expression as well as reduced protein expression (57) of several nuclear encoded subunits of complex I (27), this newly described mechanism might be involved in the induction of mitochondrial dysfunction in AD and thereby represent an exciting new pathway which could be targeted for AD therapy.

## Abbreviations

A selected list of abbreviations is available in the Supplementary Data.

## Data Availability

The data that support the findings of this study are available from the corresponding author upon request.

## Acknowledgments

We are grateful to Dr. Stephan Werner for his excellent technical assistance.

## Supplementary data

Supplementary data is uploaded as one single PDF file (attached).

## Funding

This research was funded by the Deutsche Forschungsgemeinschaft (DFG) by grants TRR319 RMaP, TP A05 and HE3397/13-2 to M.H.

## Author Contributions

Conceptualization: K.F., M.H.; biomolecular experiments: J.E.P., M.J., M.K., M.L., L.W., C.L., L.R-C., K.M.W.M., N.R., N.K., Y.M., K.E.; analysis and interpretation of the data: M.J., J.E.P, M.K., M.L., Y.M., L.R-C., N.R., F. P., W.R., N.K., A.M., S.G.; writing of the paper: M.J., J.E.P., K.F., M.H., M.L., W.R., A.M., Y.M., S.G., K.E.. KF and M.H. supervised the work. All authors have read and agreed to the published version of the manuscript.

## Conflict of Interest

The authors declare no conflict of interest.

## Notes

### Competing Interest Statement

The authors have declared no competing interest.

## References

1. Motorin, Y. and Helm, M. (2022) RNA nucleotide methylation: 2021 update. Wiley Interdiscip. Rev. RNA, 13, 1–37.

2. Freund, I., Eigenbrod, T., Helm, M. and Dalpke, A.H. (2019) RNA modifications modulate activation of innate toll-like receptors. Genes (Basel)., 10, 92.

3. Ruggieri, A., Helm, M. and Chatel-Chaix, L. (2021) An epigenetic ‘extreme makeover’: the methylation of flaviviral RNA (and beyond). RNA Biol., 18, 696–708.

4. Helm, M. and Alfonzo, J.D. (2014) Post-transcriptional RNA modifications: Playing metabolic games in a cell’s chemical legoland. Chem Biol., 21, 174–185.

5. Chen, A.Y., Owens, M.C. and Liu, K.F. (2023) Coordination of RNA modifications in the brain and beyond. Mol. Psychiatry, 10.1038/s41380-023-02083-2.

6. Boccaletto, P., Stefaniak, F., Ray, A., Cappannini, A., Mukherjee, S., Purta, E., Kurkowska, M., Shirvanizadeh, N., Destefanis, E., Groza, P., et al. (2022) MODOMICS: A database of RNA modification pathways. 2021 update. Nucleic Acids Res., 50, D231–D235.

7. Jin, H., Huo, C., Zhou, T. and Xie, S. (2022) m1A RNA Modification in Gene Expression Regulation. Genes (Basel)., 13, 910.

8. Li, X., Xiong, X., Wang, K., Wang, L., Shu, X., Ma, S. and Yi, C. (2016) Transcriptome-wide mapping reveals reversible and dynamic N1-methyladenosine methylome. Nat. Chem. Biol., 12, 311–316.

9. Grozhik, A. V., Olarerin-George, A.O., Sindelar, M., Li, X., Gross, S.S. and Jaffrey, S.R. (2019) Antibody cross-reactivity accounts for widespread appearance of m1A in 5’UTRs. Nat. Commun., 10, 1– 13.

10. Helm, M., Lyko, F. and Motorin, Y. (2019) Limited antibody specificity compromises epitranscriptomic analyses. Nat. Commun., 10, 9–11.

11. Safra, M., Sas-Chen, A., Nir, R., Winkler, R., Nachshon, A., Bar-Yaacov, D., Erlacher, M., Rossmanith, W., Stern-Ginossar, N. and Schwartz, S. (2017) The m1A landscape on cytosolic and mitochondrial mRNA at single-base resolution. Nature, 551, 251–257.

12. Li, X., Xiong, X., Zhang, M., Wang, K., Chen, Y., Zhou, J., Mao, Y., Lv, J., Yi, D., Chen, X.W., et al. (2017) Base-Resolution Mapping Reveals Distinct m 1 A Methylome in Nuclear- and Mitochondrial-Encoded Transcripts. Mol. Cell, 68, 993–1005.e9.

13. Berrisford, J.M., Baradaran, R. and Sazanov, L.A. (2016) Structure of bacterial respiratory complex i. Biochim. Biophys. Acta - Bioenerg., 1857, 892–901.

14. Vilardo, E. and Rossmanith, W. (2015) Molecular insights into HSD10 disease: Impact of SDR5C1 mutations on the human mitochondrial RNase P complex. Nucleic Acids Res., 43, 5112–5119.

15. Agip, A.N.A., Blaza, J.N., Bridges, H.R., Viscomi, C., Rawson, S., Muench, S.P. and Hirst, J. (2018) Cryo-em structures of complex i from mouse heart mitochondria in two biochemically defined states. Nat. Struct. Mol. Biol., 25.

16. Efremov, R.G., Baradaran, R. and Sazanov, L.A. (2010) The architecture of respiratory complex I. Nature, 465, 441–445.

17. Sazanov, L.A. (2015) A giant molecular proton pump: Structure and mechanism of respiratory complex I. Nat. Rev. Mol. Cell Biol., 16, 375–388.

18. Belevich, G., Knuuti, J., Verkhovsky, M.I., Wikström, M. and Verkhovskaya, M. (2011) Probing the mechanistic role of the long α-helix in subunit L of respiratory Complex I from Escherichia coli by site-directed mutagenesis. Mol. Microbiol., 82, 1086–1095.

19. Perez Ortiz, J.M. and Swerdlow, R.H. (2019) Mitochondrial dysfunction in Alzheimer’s disease: Role in pathogenesis and novel therapeutic opportunities. Br. J. Pharmacol., 176, 3489–3507.

20. Jörg, M., Plehn, J.E., Friedland, K. and Müller, W.E. (2021) Mitochondrial Dysfunction as a Causative Factor in Alzheimer’s Disease-Spectrum Disorders: Lymphocytes as a Window to the Brain. Curr. Alzheimer Res., 18, 733–752.

21. Reddy, P.H. (2007) Mitochondrial dysfunction in aging and Alzheimer’s disease: Strategies to protect neurons. Antioxidants Redox Signal., 9, 1647–1658.

22. Leuner, K., Schütt, T., Kurz, C., Eckert, S.H., Schiller, C., Occhipinti, A., Mai, S., Jendrach, M., Eckert, G.P., Kruse, S.E., et al. (2012) Mitochondrion-derived reactive oxygen species lead to enhanced amyloid beta formation. Antioxid. Redox Signal., 16, 1421–33.

23. Leuner, K., Schulz, K., Schütt, T., Pantel, J., Prvulovic, D., Rhein, V., Savaskan, E., Czech, C., Eckert, A. and Müller, W.E. (2012) Peripheral mitochondrial dysfunction in Alzheimer’s disease: focus on lymphocytes. Mol. Neurobiol., 46, 194–204.

24. Wang, X., Su, B., Lee, H.G., Li, X., Perry, G., Smith, M.A. and Zhu, X. (2009) Impaired balance of mitochondrial fission and fusion in Alzheimer’s disease. J. Neurosci., 29, 9090–9103.

25. Pradeepkiran, J.A. and Hemachandra Reddy, P. (2020) Defective mitophagy in Alzheimer’s disease. Ageing Res. Rev., 64, 1–38.

26. Grimm, A., Friedland, K. and Eckert, A. (2016) Mitochondrial dysfunction: the missing link between aging and sporadic Alzheimer’s disease. Biogerontology, 17, 281–96.

27. Holper, L., Ben-Shachar, D. and Mann, J. (2019) Multivariate meta-analyses of mitochondrial complex I and IV in major depressive disorder, bipolar disorder, schizophrenia, Alzheimer disease, and Parkinson disease. Neuropsychopharmacology, 44, 837–849.

28. Terada, T., Therriault, J., Kang, M.S.P., Savard, M., Pascoal, T.A., Lussier, F., Tissot, C., Wang, Y.T., Benedet, A., Matsudaira, T., et al. (2021) Mitochondrial complex I abnormalities is associated with tau and clinical symptoms in mild Alzheimer’s disease. Mol. Neurodegener., 16, 1–12.

29. Bobba, A., Amadoro, G., Valenti, D., Corsetti, V., Lassandro, R. and Atlante, A. (2013) Mitochondrial respiratory chain Complexes I and IV are impaired by β-amyloid via direct interaction and through Complex I-dependent ROS production, respectively. Mitochondrion, 13, 298–311.

30. Stockburger, C., Gold, V.A.M., Pallas, T., Kolesova, N., Miano, D., Leuner, K. and Müller, W.E. (2014) A cell model for the initial phase of sporadic Alzheimer’s disease. J. Alzheimers. Dis., 42, 395–411.

31. Francis, B.M., Yang, J., Song, B.J., Gupta, S., Maj, M., Bazinet, R.P., Robinson, B. and Mount, H.T.J. (2014) Reduced levels of mitochondrial complex i subunit NDUFB8 and linked complex i + III oxidoreductase activity in the TgCRND8 mouse model of Alzheimer’s disease. J. Alzheimer’s Dis., 39, 347–355.

32. Adlimoghaddam, A., Snow, W.M., Stortz, G., Perez, C., Djordjevic, J., Goertzen, A.L., Ko, J.H. and Albensi, B.C. (2019) Regional hypometabolism in the 3xTg mouse model of Alzheimer’s disease. Neurobiol. Dis., 127, 264–277.

33. Fachal, L., Mosquera-Miguel, A., Pastor, P., Ortega-Cubero, S., Lorenzo, E., Oterino-Durán, A., Toriello, M., Quintáns, B., Camiña-Tato, M., Sesar, A., et al. (2015) No evidence of association between common European mitochondrial DNA variants in Alzheimer, Parkinson, and migraine in the Spanish population. Am. J. Med. Genet. Part B Neuropsychiatr. Genet., 168, 54–65.

34. Wei, W., Keogh, M.J., Wilson, I., Coxhead, J., Ryan, S., Rollinson, S., Griffin, H., Kurzawa-Akanbi, M., Santibanez-Koref, M., Talbot, K., et al. (2017) Mitochondrial DNA point mutations and relative copy number in 1363 disease and control human brains. Acta Neuropathol. Commun., 5, 13.

35. Eckert, A., Steiner, B., Marques, C., Leutz, S., Romig, H., Haas, C. and Müller, W.E. (2001) Elevated vulnerability to oxidative stress-induced cell death and activation of caspase-3 by the Swedish amyloid precursor protein mutation.pdf. J. Neurosci. Res., 64, 183–192.

36. Keil, U., Bonert, A., Marques, C.A., Scherping, I., Weyermann, J., Strosznajder, J.B., Müller-Spahn, F., Haass, C., Czech, C., Pradier, L., et al. (2004) Amyloid β-induced changes in nitric oxide production and mitochondrial activity lead to apoptosis. J. Biol. Chem., 279, 50310–50320.

37. Holzmann, J., Frank, P., Löffler, E., Bennett, K.L., Gerner, C. and Rossmanith, W. (2008) RNase P without RNA: Identification and Functional Reconstitution of the Human Mitochondrial tRNA Processing Enzyme. Cell, 135, 462–474.

38. Stine, W.B., Jungbauer, L., Yu, C. and Ladu, M.J. (2011) Preparing synthetic Aβ in different aggregation states. Methods Mol. Biol., 670, 13–32.

39. Heiser, J.H., Schuwald, A.M., Sillani, G., Ye, L., Müller, W.E. and Leuner, K. (2013) TRPC6 channel-mediated neurite outgrowth in PC12 cells and hippocampal neurons involves activation of RAS/MEK/ERK, PI3K, and CAMKIV signaling. J. Neurochem., 127, 303–13.

40. Sims, N.R. and Anderson, M.F. (2008) Isolation of mitochondria from rat brain using Percoll density gradient centrifugation. Nat. Protoc., 3, 1228–1239.

41. Connolly, N.M.C., Theurey, P., Adam-Vizi, V., Bazan, N.G., Bernardi, P., Bolaños, J.P., Culmsee, C., Dawson, V.L., Deshmukh, M., Duchen, M.R., et al. (2018) Guidelines on experimental methods to assess mitochondrial dysfunction in cellular models of neurodegenerative diseases. Cell Death Differ., 25, 542–572.

42. Miller, J.A., Guillozet-Bongaarts, A., Gibbons, L.E., Postupna, N., Renz, A., Beller, A.E., Sunkin, S.M., Ng, L., Rose, S.E., Smith, K.A., et al. (2017) Neuropathological and transcriptomic characteristics of the aged brain. Elife, 6, 1–26.

43. Mathys, H., Davila-Velderrain, J., Peng, Z., Gao, F., Mohammadi, S., Young, J.Z., Menon, M., He, L., Abdurrob, F., Jiang, X., et al. (2019) Single-cell transcriptomic analysis of Alzheimer’s disease. Nature, 570, 332–337.

44. Montine, T.J., Sonnen, J.A., Montine, K.S., Crane, P.K. and Larson, E.B. (2012) Adult Changes in Thought study: dementia is an individually varying convergent syndrome with prevalent clinically silent diseases that may be modified by some commonly used therapeutics. Curr. Alzheimer Res., 9, 718–23.

45. Werner, S., Schmidt, L., Marchand, V., Kemmer, T., Falschlunger, C., Sednev, M. V., Bec, G., Ennifar, E., Höbartner, C., Micura, R., et al. (2020) Machine learning of reverse transcription signatures of variegated polymerases allows mapping and discrimination of methylated purines in limited transcriptomes. Nucleic Acids Res., 48, 3734–3746.

46. Kristen, M., Plehn, J., Marchand, V., Friedland, K., Motorin, Y., Helm, M. and Werner, S. (2020) Manganese ions individually alter the reverse transcription signature of modified ribonucleosides. Genes (Basel)., 11, 1–12.

47. Braak, H. and Braak, E. (1991) Neuropathological stageing of Alzheimer-related changes. Acta Neuropatholigca, 82, 239–259.

48. Chatzispyrou, I.A., Held, N.M., Mouchiroud, L., Auwerx, J. and Houtkooper, R.H. (2015) Tetracycline antibiotics impair mitochondrial function and its experimental use confounds research. Cancer Res., 75, 4446–4449.

49. Houtkooper, R.H., Mouchiroud, L., Ryu, D., Moullan, N., Katsyuba, E., Knott, G., Williams, R.W. and Auwerx, J. (2013) Mitonuclear protein imbalance as a conxerved longevity mechanism. Nature, 497, 451–457.

50. Wüst, R.C.I., Coolen, B.F., Held, N.M., Daal, M.R.R., Tazehkandi, V.A., Baks-Te Bulte, L., Wiersma, M., Kuster, D.W.D., Brundel, B.J.J.M., van Weeghel, M., et al. (2021) The antibiotic doxycycline impairs cardiac mitochondrial and contractile function. Int. J. Mol. Sci., 22, 4100.

51. Vilardo, E. and Rossmanith, W. (2013) The Amyloid-β-SDR5C1(ABAD) Interaction Does Not Mediate a Specific Inhibition of Mitochondrial RNase P. PLoS One, 8, e65609.

52. Bhatta, A., Dienemann, C., Cramer, P. and Hillen, H.S. (2021) Structural basis of RNA processing by human mitochondrial RNase P. Nat. Struct. Mol. Biol., 28, 713–723.

53. Morsy, A. and Trippier, P.C. (2019) Amyloid-Binding Alcohol Dehydrogenase (ABAD) Inhibitors for the Treatment of Alzheimer’s Disease. J. Med. Chem., 62, 4252–4264.

54. Hoernes, T.P., Heimdörfer, D., Köstner, D., Faserl, K., Nußbaumer, F., Plangger, R., Kreutz, C., Lindner, H. and Erlacher, M.D. (2019) Eukaryotic translation elongation is modulated by single natural nucleotide derivatives in the coding sequences of mRNAs. Genes (Basel)., 10, 1–12.

55. Haass, C. and Selkoe, D. (2022) If amyloid drives Alzheimer disease, why have anti-amyloid therapies not yet slowed cognitive decline? PLoS Biol., 20, e3001694.

56. McDade, E., Voytyuk, I., Aisen, P., Bateman, R.J., Carrillo, M.C., De Strooper, B., Haass, C., Reiman, E.M., Sperling, R., Tariot, P.N., et al. (2021) The case for low-level BACE1 inhibition for the prevention of Alzheimer disease. Nat. Rev. Neurol., 17, 703–714.

57. Devi, L., Prabhu, B.M., Galati, D.F., Avadhani, N.G. and Anandatheerthavarada, H.K. (2006) Accumulation of amyloid precursor protein in the mitochondrial import channels of human Alzheimer’s disease brain is associated with mitochondrial dysfunction. J. Neurosci., 26, 9057– 9068.

