## Supplementary Data for "N1-methylation of adenosine (m^1^A) in ND5 mRNA leads to complex I dysfunction in Alzheimer’s disease"

**Abbreviation list**

| Abbrevation | Description | Synonym |
|---|---|---|
| $m^1A$ | N1-methyladenosine | |
| ND5 | NADH-Dehydrogenase Subunit 5 | NADH-Ubiquinone Oxidoreductase Chain 5 |
| RNA | Ribonucleic acid | |
| mRNA | Messenger RNA | |
| tRNA | Transfer RNA | |
| (mt)tRNA | Mitochondrial tRNA | |
| (mt)mRNA | Mitochondrial mRNA | |
| rRNA | Ribosomal RNA | |
| AD | Alzheimer Disease | |
| Aβ | Amyloid β | |
| TRMT10C | TRNA Methyltransferase 10C | MRPP1; RG9MTD1 |
| SDR5C1 | Short Chain Dehydrogenase/Reductase Family 5C, Member 1 | MRPP2; ERAB; ABAD; HSD17B10; MHBD; HADH2 |
| PRORP | Protein Only RNase P Catalytic Subunit | MRPP3; KIAA0391 |
| mtRNase P | Mitochondrial RNase P | |
| ATP | Adenosine Triphosphate | |
| ROS | Reactive Oxygen Species | |
| SNP | Single Nucleotide Polymorphism | |
| RNAseq | RNA sequencing | |
| scRNAseq | Single-cell RNA sequencing | |
| HEK | Human Embryonic Kidney cells | |
| APP | Amyloid Precursor Protein | |
| APPwt | HEK 293, stabily transfected with human wild-type APP | |
| Ctl | Respective control | |
| DMEM | Dulbecco’s Modified Eagle’s Medium | |
| FCS | Fetal calf serum | |
| pTRMT10C | T-Rex-293 cells, stabily transfected with a plasmid coding for TRMT10C (tetracycline-inducable expression) | |
| FAD | Familial Alzheimer Disease | |
| Wt | Wild type | |
| TBI | Traumatic Brain Injury | |
| FPKM | Fragments per kilobase per million | |
| FC | Fold Change | |
| padj | Adjusted p-value | |
| NCBI | National Center for Biotechnology Information | |
| SRA | Sequence Read Archive | |
| NBB | Netherland Brain Bank | |
| CERAD | Consortium to Establish a Registry for Alzheimer’s Disease | |
| SD | Standard Deviation | |
| SEM | Standard Error of the Mean | |
| n.s. | Not significant | |
| p-value | Probability value | |
| PBS | Phosphate buffered saline | |
| WB | Western Blot | |

| | |
|---|---|
| RIPA | Radio-immunoprecipitation assay buffer |
| SDS | Sodium dodecyl sulfate |
| PMSF | Phenylmethylsulfonylfluorid |
| BSA | Bovine Serum Albumin |
| TBST | Tris buffered saline with Tween® |
| L | Liter |
| mL | Milliliter |
| µL | Microliter |
| µg | Microgram |
| µM | Micromolar |
| GAPDH | Glyceraldehyde-3-Phosphate Dehydrogenase |
| IB | Isolation Buffer |
| EGTA | Ethylene glycol-bis(2-aminoethylether)-N,N,N′,N′-tetraacetic acid |
| g | Gravitational force |
| min | Minutes |
| DTT | Dithiothreitol |
| SS-IV | SuperScript IV |
| RT | Reverse Transcription/ Reverse Transcriptase |
| PCR | Polymerase chain reaction |
| DNA | Deoxyribonucleic acid |
| mtDNA | Mitochondrial DNA |
| cDNA | Complementary DNA |
| BAM | Binary Alignment Map |
| PAGE | Polyacrylamide gel electrophoresis |
| h | Hours |
| oligo | Oligonucleotide |
| 6-FAM | 6-carboxyfluorescein |
| Cy5 | Cyanine-5 |
| TMRM | Tetramethylrhodamine methyl ester |
| CO2 | Carbon dioxide |
| FCCP | Carbonyl cyanide-p-trifluoromethoxyphenylhydrazone |
| OCR | Oxygen consumption rate |
| MMP | Mitochondrial membrane potential |
| ETS | Electron Transport System |
| Tetra | Tetracycline |

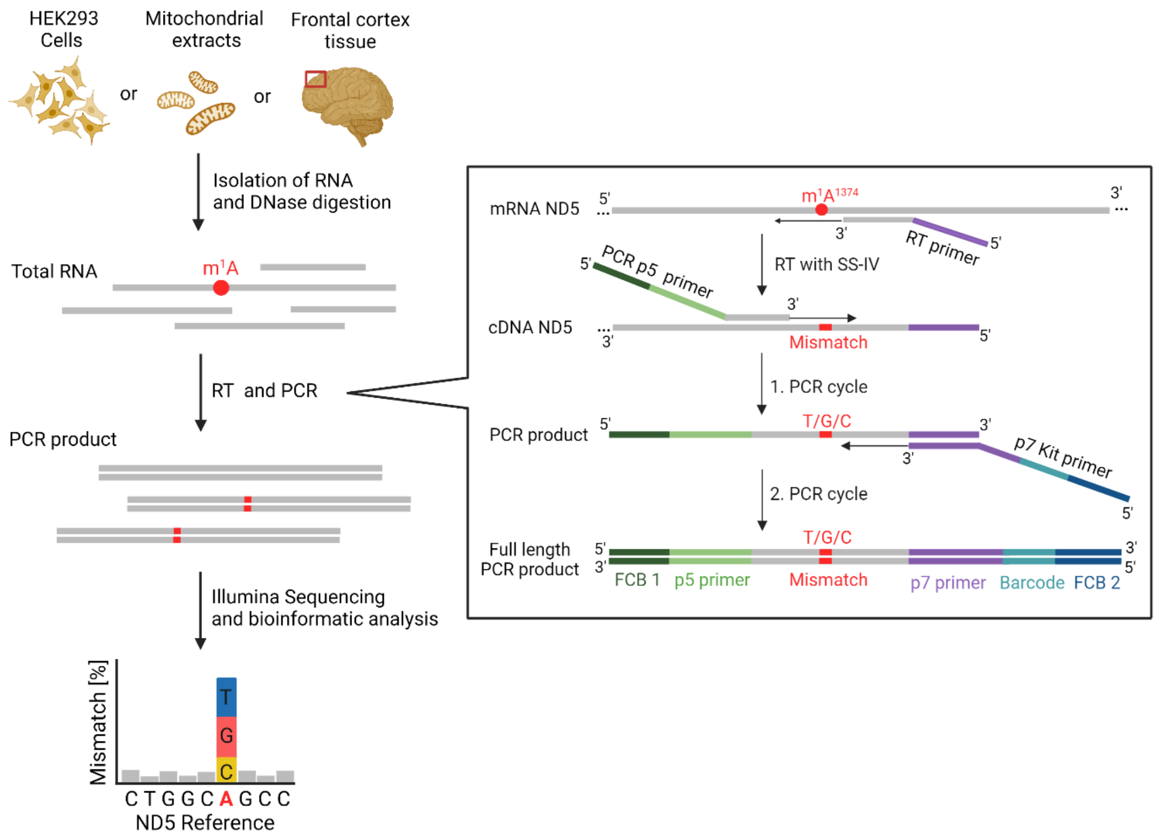

**Supplemental Figure S1| Workflow of m$^1$A analysis method.** First, RNA was isolated from HEK293 cells, mitochondrial extracts generated from HEK293 cells or frontal cortex tissue of human AD patients and healthy controls. After DNase digestion, total RNA preparations were subjected to Reverse transcription (RT) with SuperScript IV (SS-IV) and a custom-designed RT primer targeting ND5. During RT, m$^1$A is able to induce misincorporation into the newly synthesized cDNA strand at the opposite site. In the following PCR, a custom-designed PCR p5 forward primer enables polymerization of DNA complementary to the generated cDNA including T, G or C instead of A at position 1374. Next to an ND5 specific sequence, the PCR p5 primer comprises the Illumina p5 primer sequence (light green) and a flow cell binding site (FCB 1) (dark green). The PCR backward primer (p7 Kit primer) binds to a sequence which was introduced by the RT primer (purple). Besides, this p7 Kit primer contains the Illumina p7 primer sequence, a sample-specific barcode (light blue) and a second flow cell binding site (FCB 2) (dark blue). The full length PCR product was selected by gel electrophoresis (not shown) and then used for Illumina sequencing. The subsequent bioinformatic analysis yields the percentage of reads containing misincorporations at position 1374, which indicates m$^1$A$^{1374}$ methylation levels in the original sample.

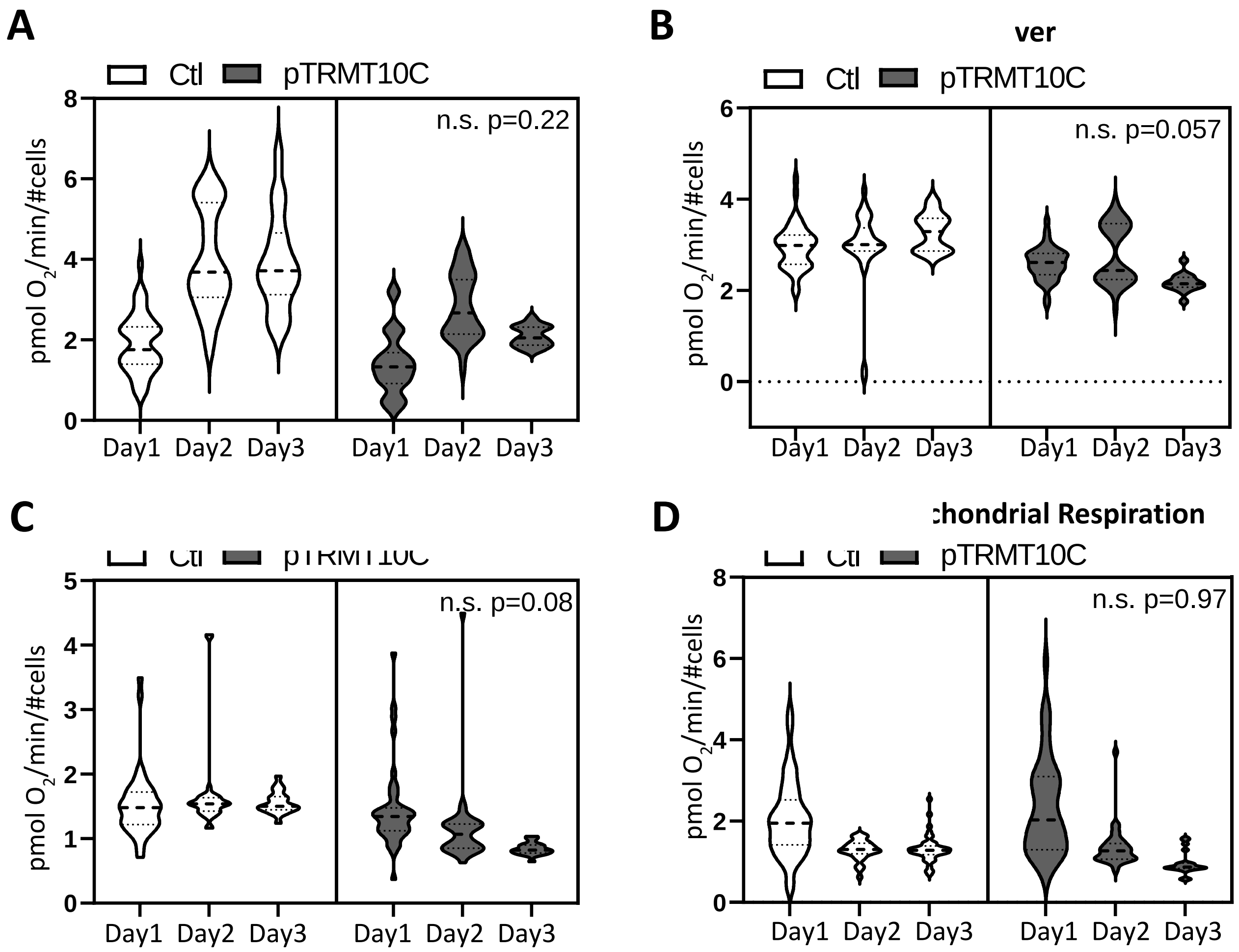

**Supplementary FIGURE S2 | TRMT10C overexpression on ETS capacitiy, ATP turnover, proton lead and non-mitochondrial respiration.** Violine plots for ETS Capacity (B), ATP Turnover (C), Proton-Leak and non-mitochondrial respiration (D) are shown from three individual experiments. n=3, nested analysis.

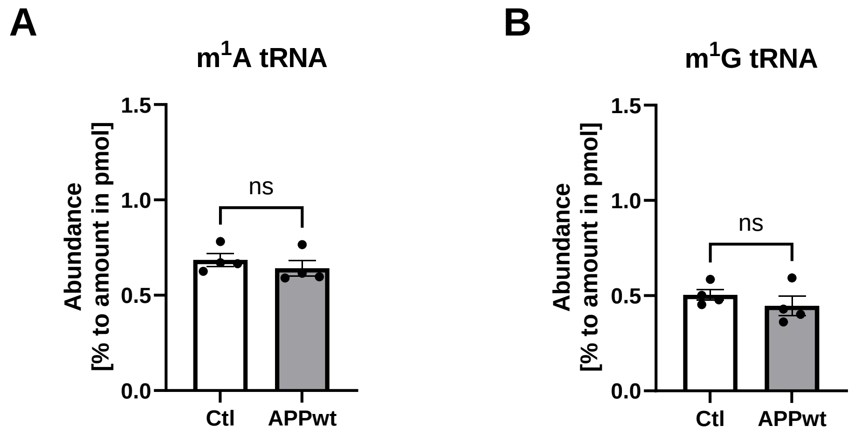

**Supplementary FIGURE S3 | m$^1$A and m$^1$G at tRNA level in HEK Ctl.** A) Abundance of m$^1$A and B) m$^1$G in HEK Ctl and HEK APPwt cells at tRNA level. No significant difference was determined. Mean ± SEM, n=4, unpaired t-test

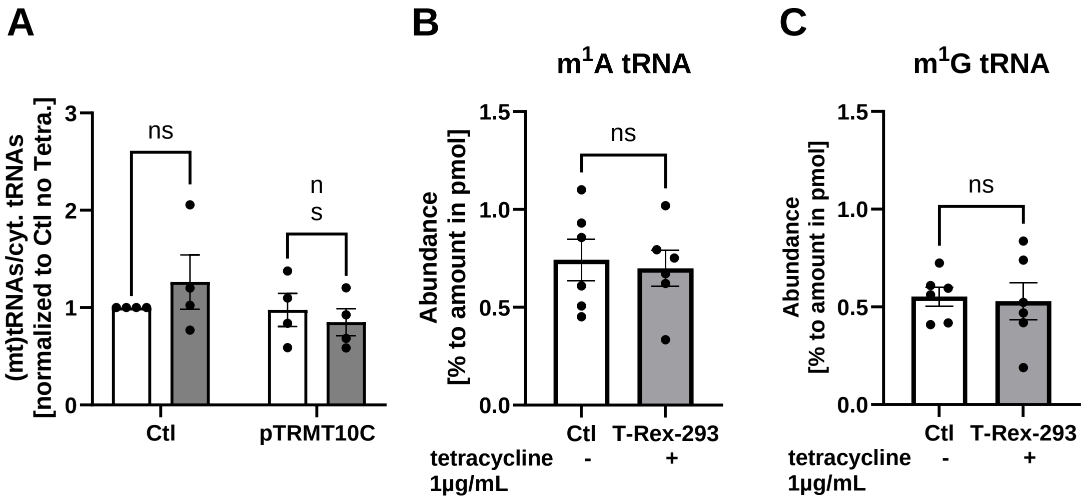

**Supplementary FIGURE S4 | m$^1$A and m$^1$G at tRNA level in tetracycline induced TRMT10C overexpressing HEK-T-Rex-293 cells.** A) The amount of mitochondrial tRNAs relative to cytosolic tRNAs is not altered in Ctl and pTRMT10C cells after treatment with 1 µg/mL Tetracycline. Mean ± SEM, n=4 biological replicates. Each biological replicate represents the mean of 3 technical replicates. B) Abundance of m$^1$A and C) m$^1$G in T-Tex-293 cells. TRMT10C expression was induced by treatment of cells with 1µg/mL of tetracycline for 24 h. Both m1A and m1G abundance analysis revealed no significant changes at tRNA level comparing TRMT10C overexpressing T-Rex-293 to non-treated T-Rex cells. Mean ± SEM, n=6, unpaired t-test.

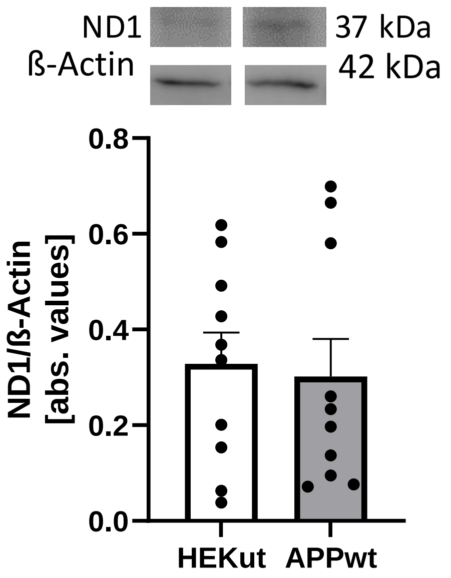

**Supplementary FIGURE S5. ND1 protein expression is not altered in HEK APPwt cells compared to HEK Ctl.** Western Blot analysis indicates no changes in HEK APPwt cells compared to HEK Ctl, n=10 Mean ± SEM, unpaired t-test.

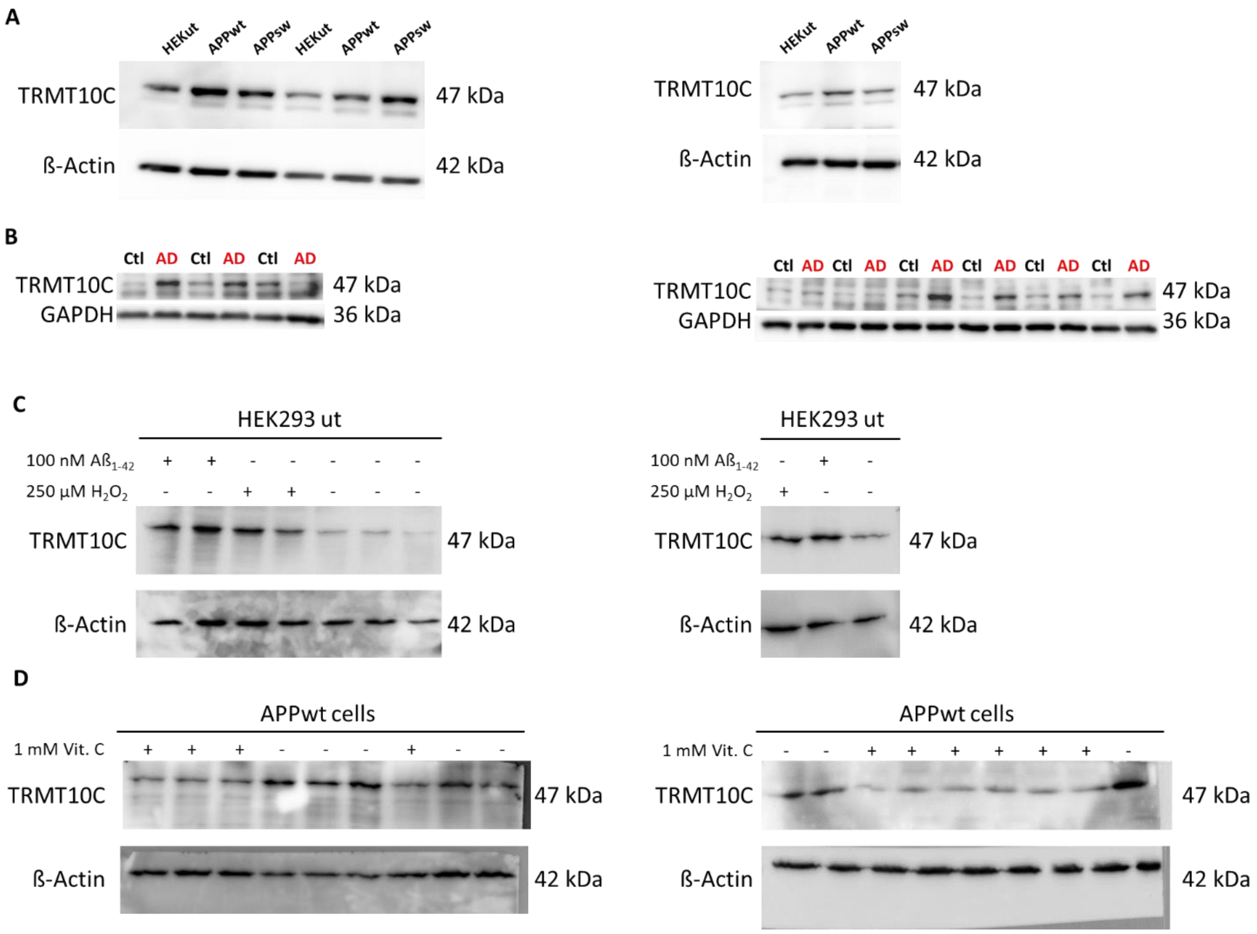

**Supplementary FIGURE S6. Representative Western Blot membranes for Figure 1.** (A) Membranes corresponding to results shown in Fig. 1C. (B) Membranes corresponding to results shown in Fig. 1D. (C) Membranes corresponding to results shown in Fig. 1G. (D) Membranes corresponding to results shown in Fig. 1H.

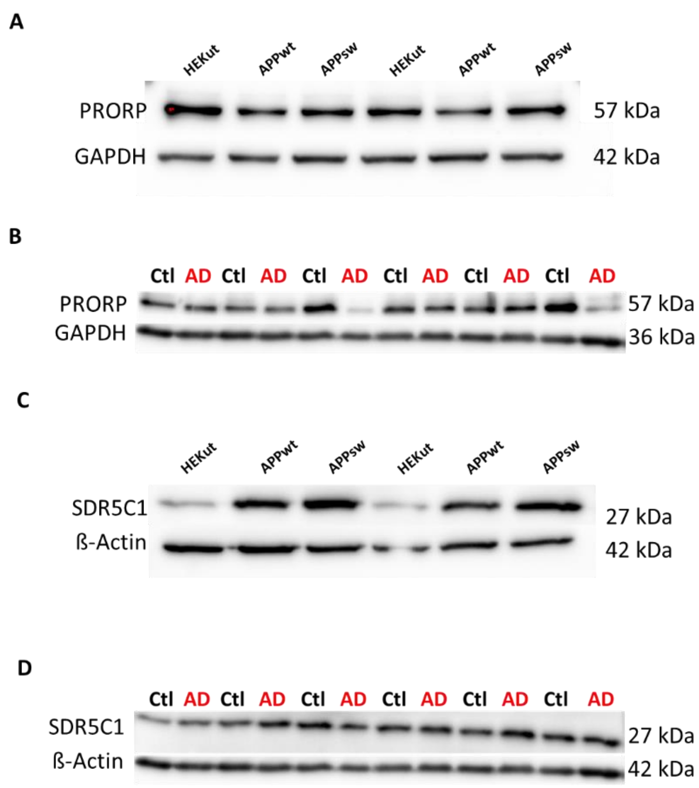

**Supplementary FIGURE S7. Representative Western Blot membranes for Figure 2.** (A) Membranes corresponding to results shown in Fig. 2B. (B) Membranes corresponding to results shown in Fig. 2C. (C) Membranes corresponding to results shown in Fig. 2E. (D) Membranes corresponding to results shown in Fig. 2F.

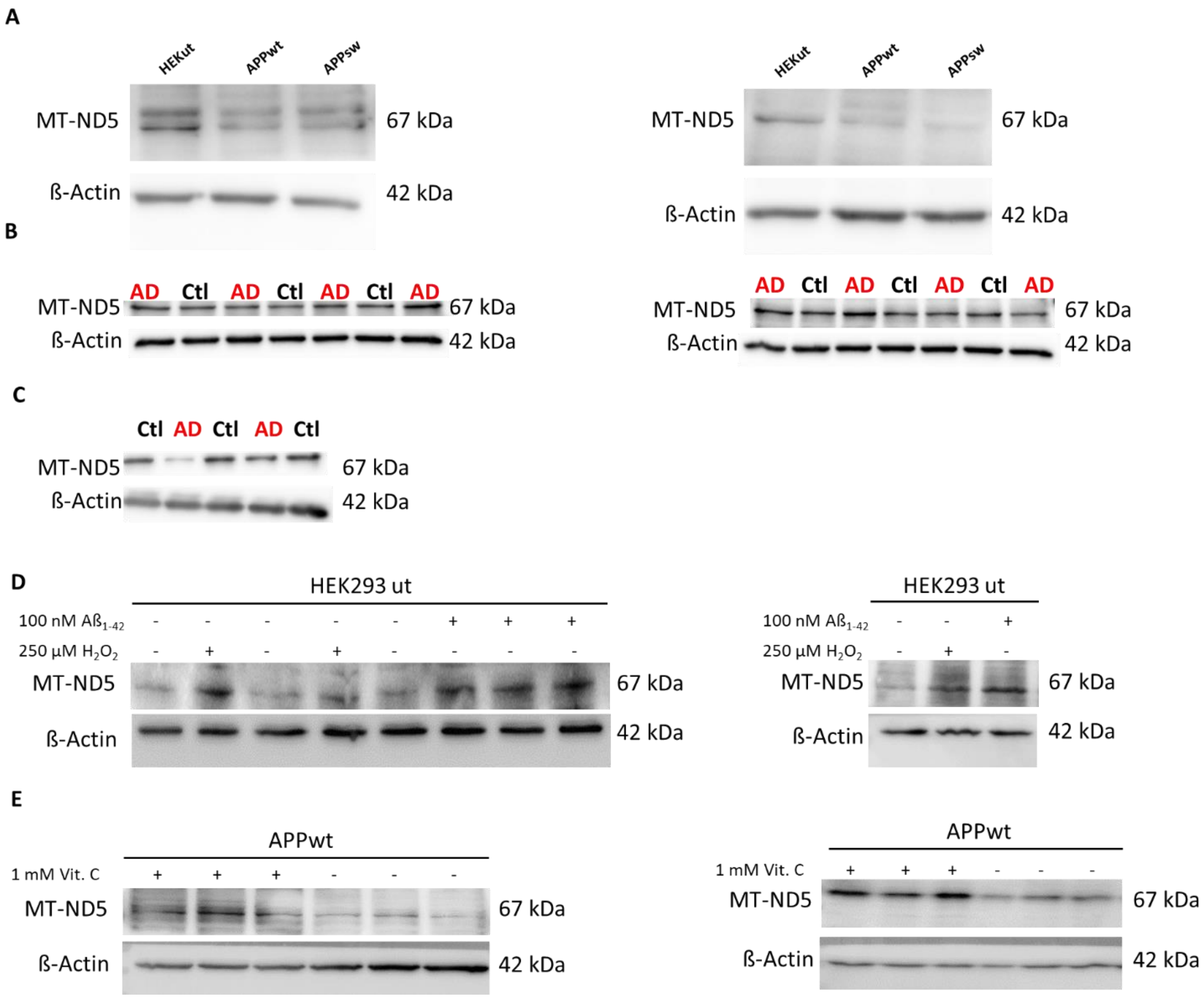

**Supplementary FIGURE S8. Representative Western Blot membranes for Figure 4.** (A) Membranes corresponding to results shown in Fig. 4A. (B) Membranes corresponding to results shown in Fig. 4B. (C) Membranes corresponding to results shown in Fig. 4C. (D) Membranes corresponding to results shown in Fig. 4H. (E) Membranes corresponding to results shown in Fig. 4I.

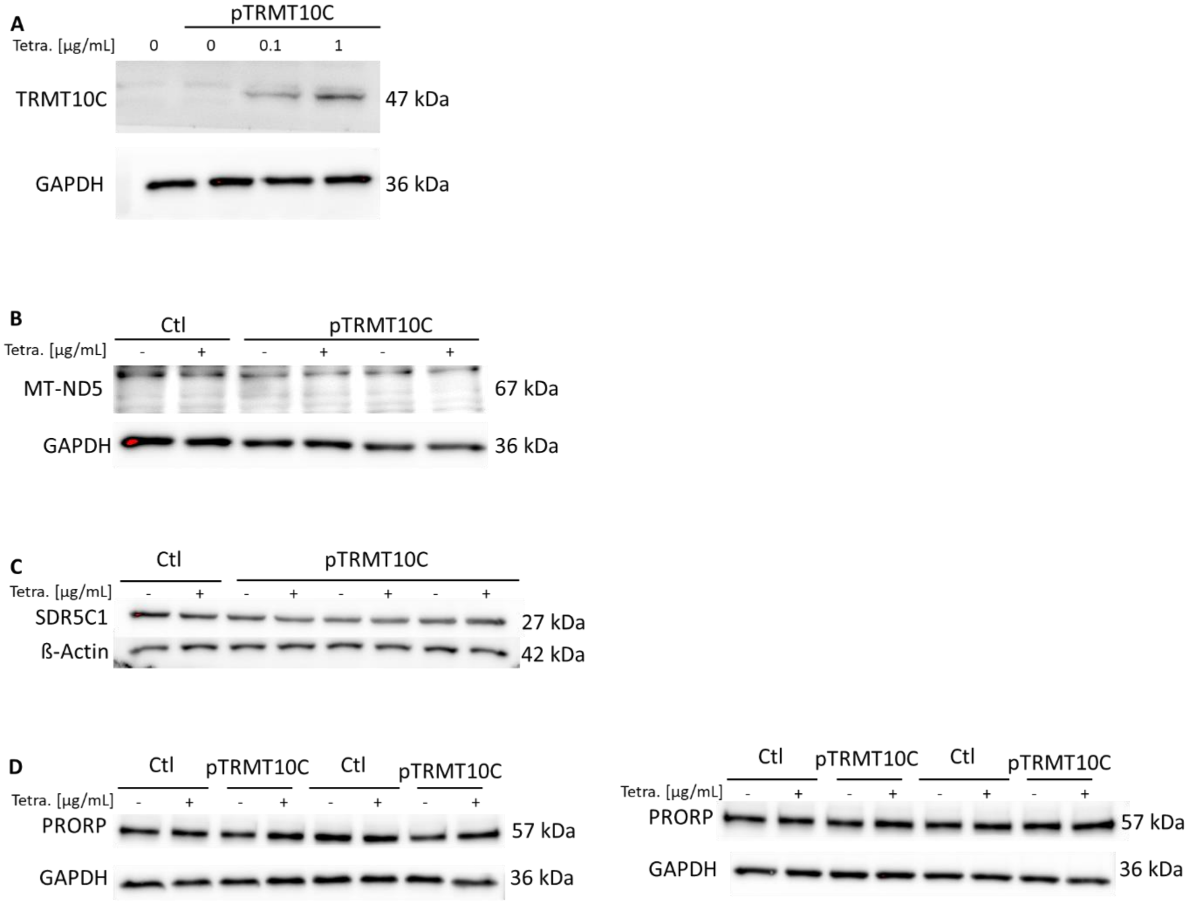

**Supplementary FIGURE S9. Representative Western Blot membranes for Figure 5.** (A) Membranes corresponding to results shown in Fig. 5A. (B) Membranes corresponding to results shown in Fig. 5C. (C) Membranes corresponding to results shown in Fig. 5D. (D) Membranes corresponding to results shown in Fig. 5E.

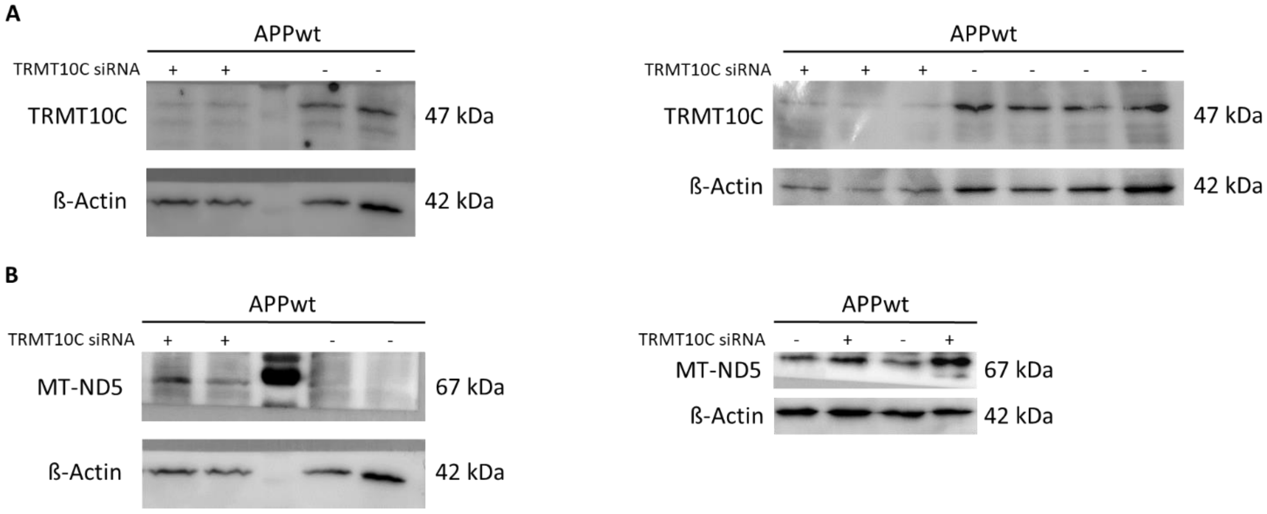

**Supplementary FIGURE S10. Representative Western Blot membranes for Figure 6.** (A) Membranes corresponding to results shown in Fig. 6A. (B) Membranes corresponding to results shown in Fig. 6D.

**Supplementary Table S1. List of primer sequences used in this study.**

| | Sequence | Additional information |
|---|---|---|
| **ND5 RT primer** | AGA CGT GTG CTC TTC CGA TCT NNN NNN NGT AAT GAG AAA TCC TGC G | NNN NNN N is a unique molecular identifier, which has not been used in the final analysis |
| **Custom P5 PCR primer** | AAT GAT ACG GCG ACC ACC GAG ATC TAC ACA CAC TCT TTC CCT ACA CGA CGC TCT TCC GAT CTA CCC CAC CCT ACT AAA CCC | |

**Supplementary Table S2. Commands, including trimming and alignment parameters.**

| |
|---|
| java -jar /path/to/Trimmomatic-0.39/trimmomatic-0.39.jar PE -phred33 <input 1> <input 2> <paired output 1> <unpaired output 1> <paired output 2> <unpaired output 2> ILLUMINACLIP:/path/to/Trimmomatic-0.39/adapters/TruSeq3-PE-2.fa:2:20:7 LEADING:30 TRAILING:30 SLIDINGWINDOW:4:20 MINLEN:8 AVGQUAL:30 |
| bowtie2 --no-1mm-upfront -D 15 -R 2 -N 0 -L 10 -i S,1,1.15 -p <number of processor cores - 2> -x '/path/to/Bowtie2_indexed_ND5_reference' -1 <mate 1> -2 <mate 2> -S <output.sam> |

**Supplementary Table S3. cDOQs for mitochondrial isolation of tRNAs.**

| tRNA | Anticodon | Sequence |
|---|---|---|
| Ala | TGC | 5'-AGACGTGTGCTCTTCCGATCTTGGTAAGGACTGCAAAACCCCACTCTGCATCAACTGAACGGATCGTCGGACTGTAGAACTCTGAAC-3' |
| Arg | TCG | 5'-AGACGTGTGCTCTTCCGATCTTGGTTGGTAAATATGATTATCATAATTTAATGAGTCGAAAGATCGTCGGACTGTAGAACTCTGAAC-3' |
| Asn | GTT | 5'-AGACGTGTGCTCTTCCGATCTTGGCTAGACCAATGGGACTTAAACCCACAAACACTTAGTTGATCGTCGGACTGTAGAACTCTGAAC-3' |
| Asp | GTC | 5'-AGACGTGTGCTCTTCCGATCTTGGTAAGATATATAGGATTTAGCCTATAATTTAACTTTGAGATCGTCGGACTGTAGAACTCTGAAC-3' |
| Cys | GCA | 5'-AGACGTGTGCTCTTCCGATCTTGGAAGCCCCGGCAGGTTTGAAGCTGCTTCTTCGAATTTGGATCGTCGGACTGTAGAACTCTGAAC-3' |
| Gln | TTG | 5'-AGACGTGTGCTCTTCCGATCTTGGCTAGGACTATGAGAATCGAACCCATCCCTGAGAATCCGATCGTCGGACTGTAGAACTCTGAAC-3' |
| Glu | TTC | 5'-AGACGTGTGCTCTTCCGATCTTGGTATTCTCGCACGGACTACAACCACGACCAATGATATGGATCGTCGGACTGTAGAACTCTGAAC-3' |
| Gly | TCC | 5'-AGACGTGTGCTCTTCCGATCTTGGTACTCTTTTTTGAATGTTGTCAAAACTAGTTAATTGGGATCGTCGGACTGTAGAACTCTGAAC-3' |
| His | GTG | 5'-AGACGTGTGCTCTTCCGATCTTGGGGTAAATAAGGGGTCGTAAGCCTCTGTTGTCAGATTCGATCGTCGGACTGTAGAACTCTGAAC-3' |
| Ile | GAT | 5'-AGACGTGTGCTCTTCCGATCTTGGTAGAAATAAGGGGGTTTAAGCTCCTATTATTTACTCTGATCGTCGGACTGTAGAACTCTGAAC-3' |
| Leu | TAG 1 | 5'-AGACGTGTGCTCTTCCGATCTTGGTACTTTTATTTGGAGTTGCACCAAAATTTTTGGGGCCGATCGTCGGACTGTAGAACTCTGAAC-3' |
| | TAA 2 | 5'-AGACGTGTGCTCTTCCGATCTTGGTGTTAAGAAGAGGAATTGAACCTCTGACTGTAAAGTTGATCGTCGGACTGTAGAACTCTGAAC-3' |
| Lys | TTT | 5'-AGACGTGTGCTCTTCCGATCTTGGTCACTGTAAAGAGGTGTTGGTTCTCTTAATCTTTAACGATCGTCGGACTGTAGAACTCTGAAC-3' |
| Met | CAT | 5'-AGACGTGTGCTCTTCCGATCTTGGTAGTACGGGAAGGGTATAACCAACATTTTCGGGGTATGATCGTCGGACTGTAGAACTCTGAAC-3' |
| Phe | GAA | 5'-AGACGTGTGCTCTTCCGATCTTGGTGTTTATGGGGTGATGTGAGCCCGTCTAAACATTTTCGATCGTCGGACTGTAGAACTCTGAAC-3' |
| Pro | TGG | 5'-AGACGTGTGCTCTTCCGATCTTGGTCAGAGAAAAAGTCTTTAACTCCACCATTAGCACCCAGATCGTCGGACTGTAGAACTCTGAAC-3' |
| Ser | GCT 1 | 5'-AGACGTGTGCTCTTCCGATCTTGGTGAGAAAGCCATGTTGTTAGACATGGGGGCATGAGTTGATCGTCGGACTGTAGAACTCTGAAC-3' |
| | TGA 2 | 5'-AGACGTGTGCTCTTCCGATCTTGGCAAAAAAGGAAGGAATCGAACCCCCCAAAGCTGGTTTGATCGTCGGACTGTAGAACTCTGAAC-3' |
| Thr | TGT | 5'-AGACGTGTGCTCTTCCGATCTTGGTGTCCTTGGAAAAAGGTTTTCATCTCCGGTTTACAAGGATCGTCGGACTGTAGAACTCTGAAC-3' |
| Trp | TCA | 5'-AGACGTGTGCTCTTCCGATCTTGGCAGAAATTAAGTATTGCAACTTACTGAGGGCTTTGAAGATCGTCGGACTGTAGAACTCTGAAC-3' |
| Tyr | GTA | 5'-AGACGTGTGCTCTTCCGATCTTGGGGTAAAAAGAGGCCTAACCCCTGTCTTTAGATTTACAGATCGTCGGACTGTAGAACTCTGAAC-3' |
| Val | TAC | 5'-AGACGTGTGCTCTTCCGATCTTGGTCAGAGCGGTCAAGTTAAGTTGAAATCTCCTAAGTGTGATCGTCGGACTGTAGAACTCTGAAC-3' |

**Supplementary Table S4. Patient details of frontal cortex samples from The Netherlands Brain Bank.**

| Individual | Diagnosis | Sex | Age | Braak stage |
|---|---|---|---|---|
| AD1 | Alzheimer's disease | f | 88 | 5 |
| AD2 | Alzheimer's disease | f | 86 | 5 |
| AD3 | Alzheimer's disease | f | 81 | 6 |
| AD4 | Alzheimer's disease | f | 88 | 4 |
| AD5 | Alzheimer's disease | f | 72 | 5 |
| AD6 | Alzheimer's disease | f | 98 | 5 |
| AD7 | Alzheimer's disease | f | 87 | 4 |
| AD8 | Alzheimer's disease | f | 76 | 4 |
| AD9 | Alzheimer's disease | m | 80 | 4 |
| AD10 | Alzheimer's disease | m | 78 | 5 |
| AD11 | Alzheimer's disease | m | 86 | 4 |
| AD12 | Alzheimer's disease | m | 97 | 5 |
| AD13 | Alzheimer's disease | m | 74 | 5 |
| Ctl1 | Non-demented control | f | 83 | 2 |
| Ctl2 | Non-demented control | f | 90 | 3 |
| Ctl3 | Non-demented control | f | 76 | 2 |
| Ctl4 | Non-demented control | f | 71 | 2 |
| Ctl5 | Non-demented control | f | 85 | 3 |
| Ctl6 | Non-demented control | f | 84 | 3 |
| Ctl7 | Non-demented control | m | 92 | 4 |
| Ctl8 | Non-demented control | m | 102 | 3 |
| Ctl9 | Non-demented control | m | 93 | 0 |
| Ctl10 | Non-demented control | m | 72 | 2 |
| Ctl11 | Non-demented control | m | 86 | 3 |
| Ctl12 | Non-demented control | m | 87 | 3 |

**Supplementary Table S5. Information about Study type, Brain region, Braak stage, Library Preparation Kit and Sex of human brain samples and human databases.**

| Dataset | Human brain tissue (NBB) | Aging, Dementia &TBI Study (aging.brain-map.org) | Mathys *et al.* (PMID: 31042697) | NCBI SRA (PRJNA720779) |
|---|---|---|---|---|
| Study type | Western Blot | RNAseq | scRNAseq | RNAseq |
| Brain region | Frontal Cortex<br>(= gyrus frontalis superior 3+4) | -Hippocampus (HIP)<br>- Parietal Cortex (PCx)<br>- White Matter of Patietal Cortex (WMPCx)<br>- Temporal cortex (TCx) | Prefrontal Cortex<br>(Brodman area 10) | - Primary visual cortex<br>- Precuneus |
| Braak stage (AD) | 4-6<br>(see Supplementary Table 1) | ⌀ 2.8 (probable AD cases, without TBI) | ⌀ 4.3 (early-stage)<br>⌀ 4.7 (all AD patients) | 6 (all samples) |
| Braak stage (Ctl) | 0-4<br>(see Supplementary Table 1) | ⌀ 2.3 (healthy controls, without TBI) | ⌀ 2.5 (all controls) | Not stated |
| Library Preperation Kit | -- | Illumina TruSeq Stranded Total RNA Sample Prep Kit (RS-122-2203) | Chromium Single Cell 3' Reagent Kits v.2 | Illumina TruSeq Stranded Total RNA Sample Prep Kit |
| Sex (AD) | 8 female + 5 male | ND5, PRORP, TRMT10C, SDR5C1<br>5-8 male / 2-3 female | 8 female + 7 male (early-stage)<br>12 female + 12 male (all AD patients) | 2 female + 3 male |
| Sex (Ctl) | 6 female + 6 male | ND5<br>12-14 male / 13-14 female<br>PRORP, TRMT10C, SDR5C1<br>13-15 male / 13-14 female | 12 female + 12 male | 2 female +3 male |
