## Supplementary Materials and Methods for "N1-methylation of adenosine (m^1^A) in ND5 mRNA leads to complex I dysfunction in Alzheimer’s disease"

### Supplementary Material and Methods

#### Cell culture

HEK 293 cells were transfected with a DNA construct harboring the human APPwt gene, inserted downstream of a cytomegalovirus promoter using the FUGENE 6 technology (Roche Diagnostics) (44, 45). The stably expressing APPwt HEK 293 cells were cultured in Dulbecco's modified Eagle's medium (DMEM) (Gibco, REF: 41965-039) supplemented with 10% heat-inactivated fetal calf serum (FCS) (Gibco, REF: 10270-106, Lot: 2156880), 50 Units/mL penicillin and 50 µg/mL streptomycin (Pen Strep, Gibco, REF: 15140-122), and 400 µg/mL G418 Solution (Roche, 4727894001) at 37 °C in a humidified incubator containing 5% $CO_2$. Control cells (Ctl) (untransfected HEK 293) were cultured in the same medium, excluding G418.

For tetracycline-induced overexpression of TRMT10C the complete coding sequence including their native initiation codon context was cloned by PCR in frame with a C-terminal FLAG-tag (46). T-Rex-293 cells (Invitrogen, R71007) were transfected and stable cell lines selected (pTRMT10C cells). Control T-Rex-293 (Ctl) cells were cultured in DMEM supplemented with 10% heat-inactivated FCS, 50 Units/mL penicillin, 50 µg/mL streptomycin and 5 µg/mL blasticidin (Thermo Fisher Scientific, R21001). Medium of pTRMT10C cells additionally contained 100 µg/mL hygromycin (Invitrogen, 10687010) for selecting the pcDNA™5/TO plasmid. 24 hours before the experimental procedure pTRMT10C as well as T-Rex-293 cells were incubated with 0 µg/mL and 1 µg/mL tetracycline (Sigma-Aldrich, T7660).

#### TRMT10C Knockdown

For siRNA-induced knockdown of TRMT10C, predesigned *Silencer*™ select siRNA against TRMT10C (Thermo Fisher Scientific, 4392420, Assay ID: s29784) and the related *Silencer*™ select siRNA negative control (Thermo Fisher Scientific, 4390843) were purchased. One day prior to transfection cells were seeded to reach a confluency of 60-80% at transfection. *Silencer*™ select siRNA against TRMT10C and negative control were diluted and mixed according to manufacturer's protocol with Lipofectamine RNAiMAX (Thermo Fisher Scientific, 13778-075) and Opti-MEM® Media (Thermo Fisher Scientific, 11058-021). Cells were incubated with TRMT10C or negative control siRNA/Lipofectamine RNAiMAX reagent for 24 hours.

#### Aß oligomerization

For analyzation of the effects of stressors on TRMT10C and ND5 expression, HEK control (ut) and APPwt cells were treated with 100 nM oligomeric Aß1-42 (AggreSure ß-Amyloid (1-42), AnaSpec, AS-72216) and 250 µM $H_2O_2$ (Sigma Aldrich®, H1009-100ML) for 24 hours. Oligomerization of $Aß_{1-42}$

was performed according to manufacturer's protocol by incubating human AggreSure ß-Amyloid (1-42) at 4°C for 24 hours. Aggregation was verified by Thioflavin T dye (AnaSpec, AS-88306) staining at 440Em/484Ex (47). Antioxidative effects were examined after treatment of APPwt cells with 1 mM Vitamin C (Sigma Aldrich®, A92902-25G) for 24 hours.

**Aß$_{1-40}$ Elisa**

Measurement of Aß1-40 levels was performed using an Amyloid beta 40 Human ELISA Kit (Invitrogen, KHB3482). Supernatants of HEKut, APPwt and siRNA treated cells (TRMT10C and negative control) cells were taken after 24 hours. The Amyloid beta 40 Human ELISA Kit assay was performed according to manufacturer's protocol.

**Preparation of protein extracts**

Cells were seeded three days in advance and synchronized with ice-cold phosphate buffered saline (PBS). 24 hours prior to use, medium was changed. For western blot (WB) analysis cells were washed with PBS twice and lysed on ice in 200 µL radio-immunoprecipitation assay buffer (RIPA) containing 50 mM Tris-HCl (pH= 7.4) (Carl Roth, 5429.5), 150 mM sodium chloride (Carl Roth, P029.3), 1% Triton™ X-100 (Carl Roth, 3051.4), 0.5% sodium deoxycholate (Sigma-Aldrich, 30970), 0.1 % sodium dodecyl sulfate (SDS) (Carl Roth, CN30.2), 5 mM ethylenediaminetetraacetic acid (Carl Roth, CN06.2) and 1 mM phenylmethylsulfonylfluorid (PMSF) (Carl Roth, 6367.1).

Mice were sacrificed by decapitation under isoflurane anesthesia. Their brains were quickly removed and three regions were dissected: hippocampus, cerebral cortex and cerebellum, and then placed in liquid nitrogen. Samples were stored at -80 °C until protein isolation with RIPA was performed.

After incubation of cell and mice samples with RIPA buffer on ice for 45 min and a centrifugation step (10 000 x g, 4°C, 10 min), proteins were located in the supernatant. Total protein content was estimated via Bradford method (Bio-Rad Protein Assay Dye Reagent Concentrate, Bio-Rad Laboratories, #5000006)).

**Western Blot**

20-40 µg protein per lane were loaded on a 10% SDS polyacrylamide gel electrophoresis (PAGE). Afterwards samples were transferred onto a polyvinylidene fluoride membrane (Thermo Scientific, REF: 88520), incubated for 1 h with 5% Bovine Serum Albumin (BSA) (Carl Roth, T844.3) blocking solution at 21 °C, washed three times with 0.5% Tris buffered saline with Tween® (TBST) (TRIS 20 mM (Carl Roth, 5429.5), sodium chloride 150 mM, 0.5% Tween® 20 (Carl Roth, 9127.1), ad 1 L water) and then incubated with primary antibodies overnight at 4 °C. Primary antibodies were purchased from

Abcam, Cambridge, United Kingdom (anti-APP, ab32136, anti-PRORP, ab185941, anti-ERAB [5F3], ab10260, anti-ND5, ab92624, anti-ND1, ab181848, anti-GAPDH, ab181602), Aviva Systems Biology, San Diego, USA (anti-TRMT10C, Arp40877_p050) and MERCK KGaA, Darmstadt, Germany (anti-β-Actin, A1978). After washing three times with 0.5% TBST, membranes were treated with horseradish peroxide conjugated secondary antibodies. For APP, PRORP, TRMT10C, ND5 and GAPDH an anti-rabbit antibody was used (Sigma-Aldrich, A0545), for ERAB and β-Actin an anti-mouse-antibody was used (Invitrogen, 31430). In the last step, membranes were washed again three times with 0.5% TBST and analyzed with Amersham ECL™ Prime Western Blotting Reagent (Cytiva, RNP2236) in Fusion Pulse TS (Vilber Lourmat, Marne-la-Vallée cedex, France). As for PRORP, Isoform 4 (57 kDa) was quantified in each Blot.

**Real-Time quantitative PCR**

Total RNA with purity values of A260/280 > 2.0 and A260/230 > 1.5 was used to synthesize single stranded cDNA by using High-Capacity cDNA Reverse Transcription Kit (Applied Biosystems™, 4368814) according to manufacturer's protocol. For qPCR Tagman® Assays for ND5 and GAPDH (Applied Biosystems™, 4331182 and 4331182) were used. Taqman® fast advanced master mix (Applied Biosystems™, 4444557) and appropriate Taqman® probes were mixed according to manufacturer's protocol. cDNA samples (1µg/20µL) were diluted to a final concentration of 5ng/µL prior before use. For each sample, triplicates of ND5 and GAPDH were prepared (3 wells for detection of target gene and 3 wells for detection of standard gene). To avoid liquid artifacts new pipette tips were used for each sample. No template controls were prepared additionally. qPCR Master Mix and samples were pipetted into appropriate wells of an optical 96-well reaction plate and covered with MicroAMP™ optical adhesive film (Applied Biosystems™, 4306311). After centrifugation (1,000 x g, 2 min, 4°C) plate was placed into QuantStudio 5 qPCR apparatus (Applied Biosystems™, USA). For detection the common qPCR Taqman® assay protocol was used according to manufacturer's protocol (50°C for 2min, 95°C for 2 min and 40 cycles 95°C for 1 second). Quality control and analysis of results was performed using the QuantStudio Design & Analysis software (V.1.4.3.) according to manufacturer's instructions.

**Generation of mitochondrial extracts**

A mitochondria isolation was done in order to enrich the desired mitochondrial mRNA. Therefor HEK cells were washed with ice-cold PBS and seeded in large cell culture plates. After two days growing, medium was changed and on the third day the isolation of mitochondria was performed based on Sims *et al.* (48).

In brief, cells were collected in ice-cold isolation buffer (IB) (320 mM Sucrose (Grüssing, 130611000U), 2 mM Ethylene glycol-bis(2-aminoethylether)-N,N,N',N'-tetraacetic acid (EGTA) (Sigma-

Aldrich, E3889), 10 mM Trizma® base (Sigma-Aldrich, T1503), pH= 7.4 at 4 °C), centrifuged (1 000 x g, 4 °C, 5 min) and resuspended in 7 mL IB containing 12%-(V/V)-Percoll® (Sigma-Aldrich, P1644). This cell suspension was homogenized using a Tissue Grinder Dounce (Wheaton, Millville, USA) by ten loose and ten tight strokes. Homogenates were carefully applied onto previously prepared discontinuous Percoll gradients (7 mL 26%-(V/V)-Percoll layered above 4 mL 40%-(V/V)-Percoll). A first ultracentrifugation step was performed in a fixed angle rotor (BECKMAN, Type 60 Ti) in a Beckman Optima™ LE-80K centrifuge (Beckman Coulter, Krefeld, Germany) (30 700 x g, 7 min, 4 °C). Afterwards lower intermediate layer was removed with a SOFT-JECT ® syringe (Hartenstein, SE33) and 17G Hamilton™ needles (Fisher Scientific, 11597784). The obtained mitochondrial extracts were united in a new tube and filled with IB up to 20 mL. After second ultracentrifugation (16 700 x g, 4 °C, 12 min) mitochondria were accumulated in a fluffy pellet. This pellet was transferred with 500 µL IB into a new tube and after a last centrifugation step (7 300 x g, 4 °C, 5 min) the resulting pellet was dissolved in 800 µL TRI Reagent® (Sigma-Aldrich, T9424) to start isolation of mitochondrial RNA. Further steps were performed as described below.

**Isolation of total and mitochondrial RNA**

Cells were seeded two days in advance and washed with ice-cold PBS for synchronization. Cells were harvested at a confluency of 70-80% in 1 mL TRI Reagent®. Dissolution of the cells was supported by vortexing and incubation at room temperature for 5 min. Next 200 µL of chloroform (Sigma-Aldrich, C2432) was added and mixed thoroughly. After a second incubation step (10 min) at 21 °C, the cell suspension was centrifuged (16 000 x g, 4 °C, 15 min). The upper aqueous phase was taken and mixed in a 1:1 ratio with isopropanol (ROTH, 7343.1) and 1 µL glycogen (Thermo Scientific, R0551) was added, incubated at 21 °C (5 min) and centrifuged (16 000 x g, 4 °C, 10 min). The RNA pellet was washed twice with ice-cold 75% ethanol including centrifugation (16 000 x g, 4 °C, 5 min). RNA was reconstituted in 10-20 µL nuclease-free water (Invitrogen, AM9930). RNA concentrations were determined using UV-VIS spectrophotometer Nanodrop 2000 (Thermo Fisher Scientific, Waltham, USA).

**Detecting misincorporation rate at position 1374 of ND5 mRNA**

*Library preparation.* Total RNA and RNA from mitochondrial extracts were incubated with DNase I (Zymo Research, E1010) and 10x DNase digestion Buffer (Zymo Research, E1010) at 21 °C (15 min) to eliminate any existing DNA contamination and a second phenol-chloroform extraction was performed as described above. For reverse transcription 2 µg RNA, 0.5 mM dNTPs (Thermo Scientific, R0192), water and a target-specific RT primer (see Supplementary Table S1) (2.5 µM) were mixed and incubated for 5 min at 75 °C and afterwards 5 min on ice. Next 10 U/µL SuperScript-IV (Invitrogen, 18090050), dithiothreitol (DTT) and 5x RT Buffer and were added, mixed and incubated for 2 hours at 50 °C. In the end, enzyme was inactivated at 80 °C for 15 min. The RT Primer contained a target-specific sequence complementary to ND5 mRNA and a primer-binding site for indexing with the NEBNext® Multiplex Small RNA Library Prep Kit for Illumina® (New England Biolabs, #E7560S) primers afterwards during polymerase chain reaction (PCR).

PCR was conducted with 0.04 U/µL Platinum SuperFi (Invitrogen, 12351010), 5x SuperFi buffer (Invitrogen, 12351010), 0.2 mM dNTPs, 5x GC Enhancer (Invitrogen, 12351010), 0.5 µM custom-designed forward primer (see Supplementary Table S1) and reverse primers of the NEBNext® Multiplex Small RNA Library Prep Kit for Illumina®. The custom p5 PCR primer contained flow cell binding sequence, sequencing primer site 1 and a target-specific sequence complementary to ND5 cDNA. The commercial i7 index primers introduced individual barcodes for each sample, contained the sequencing primer site 2 and the second flow cell binding sequence. PCR was started in Thermocycler peqSTAR (VWR Peqlab), including an initial denaturation step at 98 °C (2 min), 30 cycles at 98 °C (15 min), 63 °C (1 min) and 72 °C (2 min), and a final extension step at 72°C (5 min). PCR products were purified on a 10% denaturing polyacrylamide gel by size selection and the expected full-length product was excised. Finally, prepared libraries were quality controlled for adapter dimer formation on an Agilent 4200 TapeStation (Agilent, Santa Clara, United States). Controlled samples were then analyzed on an Illumina MiSeq platform in 2 x 75 bp paired-end mode, demultiplexed and the resulting FASTQ files used for further analysis.

*Bioinformatic analysis.* First, all FASTQ files were quality-controlled using FastQC v. 0.11.9 (Andrews, S. “FastQC”. Braham Institute) to assess presence of adapter sequences and Q-scores (49). Then reads were trimmed with Trimmomatic v. 0.39, removing any remaining Illumina-related adapter sequences. Next, the trimmed reads were aligned to the reference sequence (human MT-ND5 cDNA Sequence, Transcript-ID: ENST00000361567.2 from ensemble.org (50) using Bowtie2 v. 2.4.1 (51). The alignment output is stored in BAM format. The two BAM files, one for the forward and one for the corresponding reverse reads, were merged using SAMtools (52). Visualization of coverage and mismatches was done by IGV v. 2.8.0 (53). Mismatch and jump rate were calculated in Excel. Trimming and alignment parameters can be found in Supplementary Table S2.

**Quantification of mitochondrial and cytosolic tRNAs**

Relative content of mitochondrial tRNAs compared to the cytosolic tRNA pool was quantified in pTRMT10C and control cells with and without addition of tetracycline (24 h, 1 µg/mL). Therefore, fluorescently labeled probes comprising a 40 nt DNA stretch complementary to the tRNA 3’ end of each mitochondrial and cytosolic tRNA isoacceptor were custom-designed (purchased at biomers.net). Probes targeting the mitochondrial fraction were Cyanine-5 (Cy5) labeled, with 6-carboxyfluorescein (6-FAM) label on hybridization oligonucleotides (oligos) targeting the cytosolic tRNA pool. A mixture of 22 mitochondrial and 44 cytosolic oligos was hybridized with total RNA and afterwards run on a native PAGE gel to physically separate oligo-tRNA hybrids from excess oligo. Finally, fluorescence intensities of hybrid bands were measured and the ratio of Cy5 (mitochondrial tRNAs) to 6-FAM (cytosolic tRNAs) compared across cell lines and biological replicates.

**Selective isolation of mitochondrial tRNAs from total HEK cell RNA**

Total HEK-cell RNA was hybridized to an equimolar mixture of cDNA oligos for quantification (cDOQs, biomers) with an amount of 20 fmol of each cDOQ per µg of total HEK-cell RNA. The cDOQ were tagged with a 5'-Cy5 fluorescence tag and the respective sequences are listed in (Supplementary Table S3). The hybridization was carried out in 30 mM HEPES (pH 7.5, Carl Roth) and 100 mM potassium acetate (Carl Roth) with a thermocycler (PEQLAB) using the gradient 95 °C (2 min), 90 °C (1 min), 85 °C (1 min) , 80 °C (1 min) , 75 °C (1 min) , 70 °C (1 min) , 60 °C (3 min) , 50 °C (3 min) , 40 °C (3 min) , 30 °C (5 min) , 25 °C (5 min), 4 °C (∞). tRNA-cDNA duplexes were isolated from unbound RNAs and cDOQs via native polyacrylamide gel electrophoresis (PAGE) with an acrylamide concentration of 10% according to their size. Gel ingredients were obtained from Carl Roth. Gel pieces containing the respective bands were cut out and resuspended in 300 µL 0.5 M ammonium acetate solution (Merck). The mixture was frozen at -80 °C for 15 minutes and shaked over night at 15 °C and 700 rpm in a thermoshaker mixer (Starlab). The liquid supernatant and the gel pieces were transferred to a Nanosep spin column (Pall) and centrifuged at 2000 g (6000 rpm) for 5 min. 1 µL glycogen (20 $\frac{\mathrm{mg}}{\mathrm{mL}}$, Thermo Fischer) and 700 µL ethanol (100%, Carl Roth) with a temperature of -80°C were added to the filtrate. The solution was vortexed, incubated for one hour at -80°C and then centrifuged for one and a half hour at 4°C and 18000 g (13000 rpm). The supernatant was removed. The nucleic acid pellet was washed with 300 µL ethanol (75%) and centrifuged for 30 min at 4°C and 18000 g (13000 rpm). The supernatant was discarded, and the pellet was dried on ambient conditions. The pellet was resuspended in MP-$H_2O$.

**Quantification of mitochondrial tRNA modifications**

160 ng of isolated tRNA-DNA hybrids per sample was digested to nucleosides using 0.6 U nuclease P1 from *P. citrinum* (Sigma-Aldrich), 0.2 U snake venom phosphodiesterase from *C. adamanteus* (Worthington), 0.2 U bovine intestine phosphatase (Sigma-Aldrich), 10 U benzonase (Sigma-Aldrich), 200 ng Pentostatin (Sigma-Aldrich) and 500 ng Tetrahydrouridine (Merck-Millipore) in 5 mM Tris (pH 8) and 1 mM magnesium chloride for two hours at 37°C. 135 ng of isolated tRNA-DNA hybrids was mixed with 25 ng of internal standard ($^{13}C$ stable isotope-labeled nucleosides from *S. cerevisiae*) and subjected to LC-MS/MS analysis (Agilent 1260 Infinity system in combination with an Agilent 6470 Triple Quadrupole mass spectrometer equipped with an electrospray ion source). The solvents consisted of 5 mM ammonium acetate buffer (pH 5.4, adjusted with acetic acid; solvent A) and LC-MS grade acetonitrile (solvent B; Honeywell). A C18 reverse HPLC column (Synergi™ 4 µM particle size, 80 Å pore size, 250 × 2.0 mm; Phenomenex) was used at a temperature of 35°C. The compounds were eluted with a constant flow rate of 0.35 mL/min and a linear gradient of 0-8% solvent

B over 10 min, followed by 8-40% solvent B over 10 min was applied. Initial conditions were regenerated with 100% solvent A for 10 min. Adenosine was detected photometrically at 254 nm via a diode array detector (DAD). The following mass parameters were used: gas temperature 300°C, gas flow 7 L/min, nebulizer pressure 60 psi, sheath gas temperature 400°C, sheath gas flow 12 L/min, capillary voltage 3000 V, nozzle voltage 0 V. The MS was operated in the positive ion mode using Agilent MassHunter software in the dynamic MRM (multiple reaction monitoring) mode. For absolute quantification, external calibration was applied as described in Thüring et al. (2016).

**Mitochondrial membrane potential**

Mitochondrial membrane potential was analyzed in HEK pTRMT10C and corresponding control cells treated and untreated with 1 µg/mL tetracycline for 24 h. 10 000 cells were seeded in a Greiner black 96-well plate with a transparent bottom. On the experiment day, the medium was changed and replaced with Tyrode´s buffer (137 mM NaCl, 2.7 mM KCl, 1 mM MgCl2, 1.8 mM CaCl2, 0.2 mM Na2HPO4, 12 mM NaHCO3, 5.5 mM D-glucose) containing tetramethylrhodamine, methyl ester (TMRM, Sigma-Aldrich, T5428) 25 nM for 15 min at 37 °C to evaluate the mitochondrial membrane potential. Nuclei were stained with 1 µg/mL Hoechst (Thermo Scientific, H1399. Live cell imaging was performed with the dyes present during image acquisition at 37 °C and 5% CO2 with a 40x magnification in an automated confocal spinning disk high-content screening system (Opera Phenix™, Perkin Elmer, Waltham, MA, USA). Z-stacks with 6 planes (0.5- 1µm) and 24 images per well were acquired. Carbonyl cyanide-p-trifluoromethoxyphenylhydrazone (FCCP, Sigma-Aldrich, C2920) 2 µM was added after imaging to quench the TMRM signal. Quantitative image analysis was done with the PhenoLOGICTM machine learning algorithm.

**Respiration assay**

Respiration was assessed using the Seahorse XF Cell Mito Stress Kit 103015-100 (Agilent Technologies, Santa Clara, CA, USA). Oxygen consumption rate (OCR) was measured in an XF96 Extracellular Flux analyzer (Seahorse Bioscience, MA USA). 80 000 cells were seeded per well one day before the experiment on Seahorse XF-96 plates previously coated with Poly-L-lysine (Sigma-Aldrich, P4707). On the day of the assay, cells were washed once with PBS, the growth medium was replaced for XF Base Medium supplemented with 1 mM pyruvate, 2.5 mM glucose, and 2 mM L-glutamine and incubated for 1 h in a non-CO2 incubator at 37°C. The measurements of basal OCR were recorded 3 times for 12 min before each injection. 1 µM of Oligomycin (Sigma-Aldrich, O4876, ATP synthase inhibitor), 1µM pf FCCP (mitochondrial uncoupler), and a mixture of 1µM of Rotenone (Sigma-Aldrich, R8875)/Antimycin (Sigma-Aldrich, A8674, inhibitors of the complexes I and III of the electron transport chain, respectively) were injected sequentially, and three measurements were taken during 21 min.

During the last injection, cells were labeled with Hoechst 1.7 µM to quantify the number of nuclei per well for further data normalization.

Mitochondrial respiration was also evaluated using the OROBOROS Oxygraph-2k high-resolution respirometer. Briefly, two million cells were seeded two days before the experiment in 10 $cm^2$ dishes. 24 h later, the regular growth media was replaced with DMEM containing 1 µg/mL tetracycline. The next day, cells were trypsinized, counted, and 2 million cells were added per chamber. The cellular routine respiratory state, the mitochondrial coupling state, non-coupled respiratory capacity, and rotenone-antimycin or residual oxygen consumption were tested using 2 µM of Oligomycin, 1 µM of FCCP, and 1 µM of Rotenone and Antimycin.

**Human databases**

The data for the Aging, Dementia and Traumatic Brain Injury (TBI) Study (primary publication: Miller J. A., *et al.* (54)) was retrieved from http://aging.brain-map.org/download/index on the 20th of January 2020.

The normalized and non-normalized FPKM (fragments per kilobase per million) values for the genes SDR5C1, TRMT10C, MT-ND5 and PRORP were extracted from the dataset and saved in an Excel sheet (.xslx file) together with the associated donor information using a custom R script (R Core Team 2018, 3.6.0, “Planting of a Tree”). For our analysis only normalized FPKM values, corrected for RIN and batch effects, were used. Unfortunately, only 2 female patients were available for analysis. Therefore, male patients with a confirmed AD diagnosis and without TBI incidents were selected for analysis, and normalized FPKM values were compared with healthy controls.

For single-cell transcriptomic analysis, fold change (FC) and adjusted p-values (padj) of TRMT10C and SDR5C1 were used for six major brain cell types isolated from prefrontal cortex, namely excitatory neurons, inhibitory neurons, astrocytes, oligodendrocytes, precursor-oligodendrocytes and microglia from Mathys *et al.* (55) . We visualized the FC and padj of the comparison early-stage AD vs. controls (Ctl) and all AD vs. Ctl as a heatmap using the python package *seaborn* (56). Early-stage AD subjects were classified based on having amyloid burden, but modest neurofibrillary tangles and cognitive impairment.

**Human brain samples**

Human brain material was provided via the rapid autopsy program of the Netherlands Brain Bank (NBB), which provides post-mortem specimens from clinically well documented and neuropathologically confirmed cases. All cases were neuropathologically confirmed, using conventional biochemical techniques, and diagnosis performed using the Consortium to Establish a

Registry for Alzheimer's Disease (CERAD) criteria. Non-disease controls had no history or symptoms of neurologic or psychiatric disorders and were clinically no demented. Gyrus frontalis superior 3+4 was selected for controls as well as AD patients (more information in Table 1. and Supplementary Tables S4 and S5).
